# Structural basis of TMPRSS2 zymogen activation and recognition by the HKU1 seasonal coronavirus

**DOI:** 10.1101/2024.02.21.581378

**Authors:** Ignacio Fernández, Nell Saunders, Stéphane Duquerroy, William H. Bolland, Atousa Arbabian, Eduard Baquero Salazar, Catherine Blanc, Pierre Lafaye, Ahmed Haouz, Julian Buchrieser, Olivier Schwartz, Félix A. Rey

**Affiliations:** Institut Pasteur, Université de Paris Cité, CNRS UMR 3569, Unité de Virologie Structurale, 75015, Paris, France; Institut Pasteur, Université de Paris Cité, CNRS UMR 3569, Unité de Virus & Immunité, 75015, Paris, France; Institut Pasteur, Université de Paris Cité, INSERM U1222, Nanoimaging core, 75015, Paris, France; Institut Pasteur, Université de Paris Cité, Pasteur-TheraVectys Joint Lab, Paris, France; Institut Pasteur, Université Paris Cité, CNRS UMR 3528, Platform d’ingénierie des anticorps-C2RT, 75015, Paris, France; Institut Pasteur, Université Paris Cité, CNRS UMR 3528, Plateforme de cristallographie-C2RT, 75015, Paris, France

## Abstract

The human seasonal coronavirus HKU1-CoV, which causes common colds worldwide, relies on the sequential binding to a cell-surface glycan and to TMPRSS2 for entry into target cells. TMPRSS2 is a cell surface protease synthesized as a zymogen that undergoes autolytic activation to process its substrates. Several respiratory viruses - in particular coronaviruses - use TMPRSS2 for proteolytic priming of their surface spike protein to drive membrane fusion upon receptor binding. We describe the crystal structure of the HKU1-CoV receptor binding domain in complex with TMPRSS2, showing that it recognizes residues lining the catalytic groove. Combined mutagenesis of interface residues and comparison across species highlight positions 417 and 469 as determinants of HKU1-CoV host tropism. The structure of a receptor-blocking nanobody in complex with zymogen or activated TMPRSS2 further provides the structural basis of the TMPRSS2 activating conformational change, altering loops recognized by HKU1-CoV and dramatically increasing its binding affinity.

## INTRODUCTION

The cell surface transmembrane serine protease 2 (TMPRSS2), which proteolytically primes the spike protein of multiple coronaviruses for entry into cells of the respiratory tract, was recently identified as the entry receptor of human HKU1-CoV ^1^. TMPRSS2 belongs to the “type 2 transmembrane serine proteases” (TTSP) involved in proteolytic remodeling of the extracellular matrix. Dysregulation of TMPRSS2 and other TTSPs has been observed in malignancies and is associated to tumor proliferation and invasiveness ^2–5^. Moreover, TMPRSS2 is an androgen regulated protease that activates proteolytic cascades promoting prostate cancer metastases ^6^.

HKU1-CoV is a seasonal beta-coronavirus causing common colds worldwide, with complications in young children, the elderly and immunocompromised individuals ^7^. Entry of HKU1-CoV into cells relies on its trimeric spike surface glycoprotein, which catalyzes the fusion of the viral envelope with the target cell membrane. The spike is synthetized as a precursor that is cleaved into two subunits, S1 and S2. S1 harbors an N-terminal domain (NTD), a receptor binding domain (RBD), and subdomains SD1 and SD2. The RBD adopts conformations known as ‘up’ and ‘down’, which determine different forms on the spike (closed, 1-RBD-up, 2-RBD-up, 3-RBD-up or “open”). The S2 subunit is the membrane fusion effector and requires further cleavage at a second site - the S2’ site, to be functional ^8^. As an important entry factor, one of the roles of TMPRSS2 is to cleave HKU1-CoV S2 at the S2’ site ^1^, priming it for driving membrane fusion.

The HKU1-CoV spike NTD binds α2,8-linked 9-O-acetylated disialosides (9-O-Ac-Sia(α2,8)Sia) ^9–11^ causing a conformational change that exposes the RBD and opens the spike ^12^. The transition into open states has been associated to the activation of the protein to trigger membrane fusion ^13,14^. The surface sialo glycan is thus a primary receptor of HKU1-CoV, and its recognition is a specific cue for spike opening, allowing the interaction with a protein receptor ^12^. We showed that TMPRSS2 serves as a functional proteinaceous receptor for HKU1-CoV by recognizing the RBD in a so far uncharacterized mode ^1^. Even though we showed that sialic acid is important to trigger efficient HKU1-CoV entry into TMPRSS2 expressing cells ^1^, no experimental evidence has demonstrated the interplay between the two receptors. We also developed nanobodies against TMPRSS2, some of which bind without affecting recognition of the RBD (i.e., nanobody A01), while others block the interaction, reducing infection of HKU1-CoV susceptible cells, and inhibit the TMPRSS2 proteolytic activity (i.e., nanobody A07) ^1^.

TMPRSS2 is a type II single-pass transmembrane protein with an N-terminal cytosolic tail and an ectodomain containing a low-density lipoprotein receptor type-A (LDLR-A) domain followed by a class A scavenger receptor cysteine-rich (SRCR) domain and a C-terminal trypsin-like serine peptidase (SP) domain ^15^. It is synthesized as a proenzyme or zymogen that undergoes auto cleavage to reach its mature, active conformation ^16,17^. The active site of the enzyme contains the catalytic triad H296-D345-S441, which cleaves after arginine 255 of the zymogen, at the peptide sequence (RQSR_255_τIVGG), where the arrow denotes the scissile bond. The substrate sequence, designated P4-P3-P2-P1τP1’-P2’-P3’-P4’, is recognized through specific TMPRSS2 residues that form the corresponding S4-S3-S2-S1-S1’-S2’-S3’-S4’ sites along the catalytic groove ^18^. TMPRSS2 substrates include proteins of the tumor microenvironment ^6^, such as the hepatocyte growth factor (cleaved sequence KQLR_494_τVVNG), tissue plasminogen activator (PQFR_310_τIKGG), human glandular kallikrein 2 (IQSR_24_τIVGG) as well as the S2’ site of the coronavirus spike ^19–23^, at the sequences SSSR_900_τSLLE (HKU1-CoV isolate N5) or PSKRτSFIE (SARS-CoV-2 Wuhan). The amino acid cleavage preferences at positions P1-P4 of the substrate have been determined by positional scanning of synthetic peptide libraries ^6^.

Here, we determined the X-ray structure of TMPRSS2 in its zymogen form as a ternary complex with the HKU1-CoV genotype B RBD (from here on, HKU1B RBD) and the non-competing nanobody A01. The structure indicated that TMPRSS2 binds exclusively to RBDs in the ‘up’ conformation, making spike opening mandatory for binding. We provide experimental evidence supporting that sialo glycan binding to the spike at the NTD is a required first step that allows fusion with TMPRSS2 expressing cells. We further identified key TMPRSS2 residues determining host tropism through extensive functional validation of the RBD-TMPRSS2 interface. We also determined a high-resolution X-ray structure of TMPRSS2 in complex with nanobody A07, showing that it inserts its long CDR3 into the protease’s active site, explaining its role in blocking HKU1-CoV entry and TMPRSS2 activity. The crystals of the TMPRSS2/A07 complex contained two forms, the zymogen and the activated form of the protease, illustrating the activation mechanism of TMPRSS2 and providing key information for the development of specific, zymogen-targeting drugs. We also found that the conformational change caused by the activating cleavage of TMPRSS2 involves loops that are recognized by the HKU1-CoV RBD, and we show that protease activation is required for high-affinity binding.

## RESULTS

### The HKU1B RBD partially blocks the TMPRSS2 catalytic groove

We determined the X-ray structure of the ternary TMPRSS2^S441A^/RBD/A01 complex (Figure SI1) to a 3.5 Å resolution, displayed in Figure 1A-E, with the crystallographic statistics listed in Table 1. We used the inactive TMPRSS2^S441A^ mutant to increase the protein yield and avoid protein degradation during purification and crystallization. As described previously, the SP domain of TMPRSS2 has a characteristic chymotrypsin like fold ^18^, featuring two 𝛽-barrels with the substrate binding groove in between ^24^. This groove is surrounded by eight loops: loops 1, 2 and 3 in the C-terminal 𝛽-barrel controlling specificity at the P1 position of the substrate, and loops A, B, C and E, (N-terminal 𝛽-barrel) as well as loop D (C-terminal 𝛽-barrel), which affect specificity at more distal positions ^25,26^. The structure of the complex (Figure 1B) shows that the elongated HKU1B RBD has a structure resembling a pincer plier at its distal end (in the “insertion” domain, Figure 1A), with two jaws (j1 and j2) that grab the TMPRSS2 SP domain at a surface involving loops 1-3 (L1-3) as well as loop C (LC). The RBD pincer plier’s base is made by a distal 𝛽-hairpin of the RBD (residues 503-519, 𝛽10-𝛽11), with the loop connecting the two 𝛽-strands forming one of the jaws (j1, aa 509-512) while the other is formed by the segment immediately downstream the 𝛽-hairpin (j2, residues 519-533) (Figure 1B). The base of the plier contacts loops 2 and 3 of the SP domain, the j1 jaw inserts in the groove and makes contacts loops 3 and C, and j2 interacts with loop 1 (Figure 1B). The RBD thus interacts with both 𝛽-barrels of the TMPRSS2 SP domain, at either side of the groove between them, thereby obstructing access of the substrate to the catalytic site. The nanobody A01 binds at the interface between the SCRC and SP domains, and contacts loops of the SP C-terminal 𝛽-barrel at the side opposite to the catalytic groove (Figure SI1B), with a large, buried surface area (BSA) of close to 2500 Å^2^ (1300 Å^2^ on the nanobody and 1200 Å^2^ on TMPRSS2) (Table 2).

**Figure 1.**
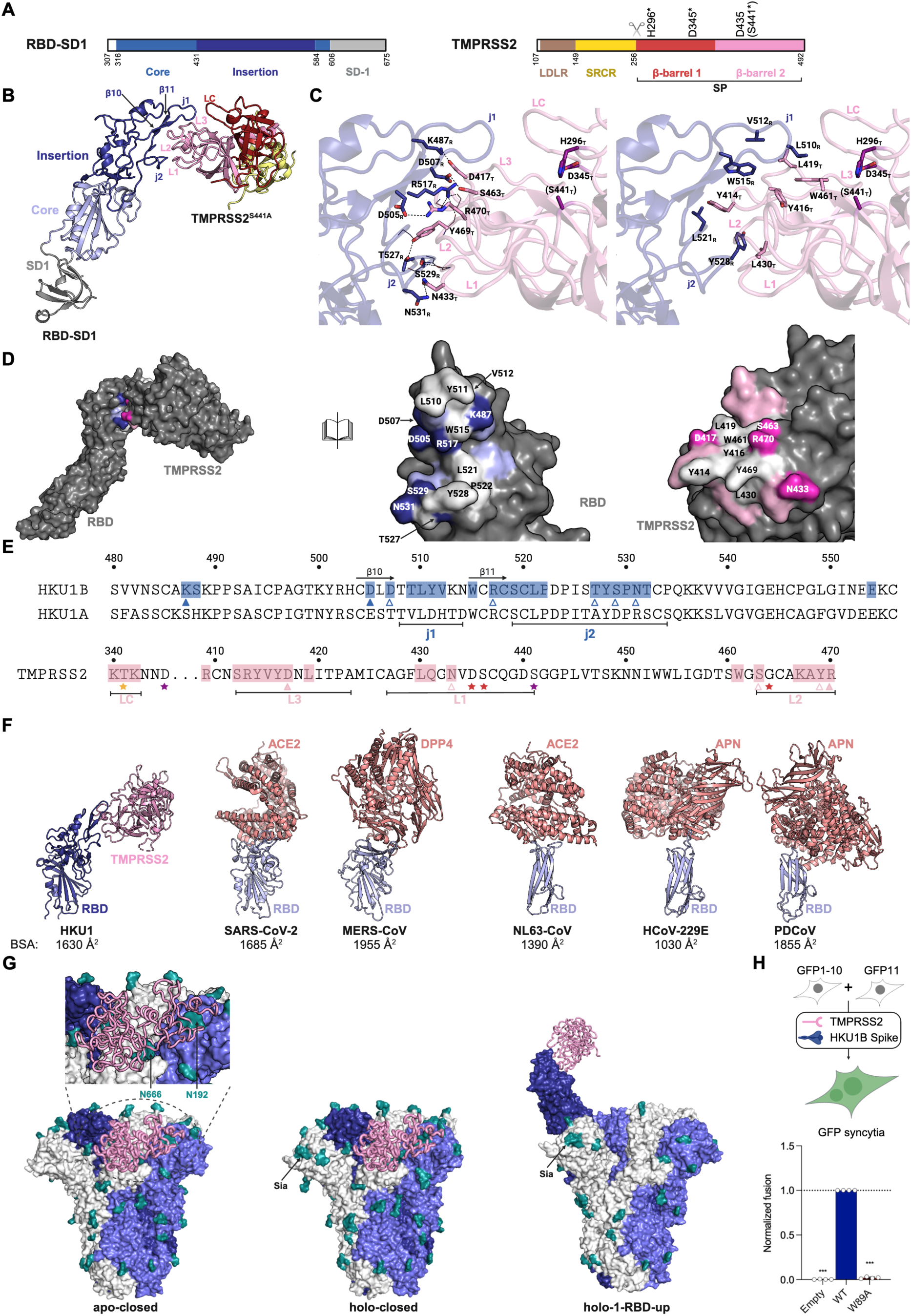
The HKU1B-RBD recognizes TMPRSS2 with a unique binding mode. (A) Schematic representation of the HKU1B SD1-RBD and TMPRSS2 constructs used for crystallization experiments. The different (sub)domains are indicated with colors, and the residues at the boundaries are indicated below the scheme. TMPRSS2 serine-peptidase (SP) domain is formed by two β-barrels, which are shown in different colors. Important TMPRSS2 residues, such as H296-D345-S441 (catalytic triad, indicated with a star) and D435 (part of the S1 site) are on top of the scheme. (S441) indicates that this active site residue was mutated to alanine in the crystallized structure. The scissor indicates the TMPRSS2 autocleavage site. (B) Crystal structure of the HKU1B-RBD in complex with TMPRSS2^S441A^. Crystals were obtained for the ternary complex with VHH-A01 but for better clarity the nanobody is not displayed (the structure of the ternary complex is displayed in Figure SI 1B). Both proteins are colored according to the (sub)domain code presented in panel (A). Important elements on the RBD, such as β-strands β10 and β11 of the pincer plier motif, and the loops that form jaws 1 (j1) and 2 (j2) are indicated. The TMPRSS2 loops at the interface are labeled L1 (loop 1, residues 427-441), L2 (loop 2, residues 462-471), L3 (loop 3, residues 412-423) and LC (loop C, residues 334-346). (C) Description of relevant contacts at the interface between the RBD and TMPRSS2^S441A^. The left panel shows polar residues that form salt bridges or hydrogen bonds (dashed lines), while the right panel shows hydrophobic and aromatic residues. The subscript of each residue indicates if it is part of the RBD (R) or TMPRSS2 (T). The residues that form the catalytic triad (H296, D345, S441A) are colored in purple. (S441) indicates that this active site residue was mutated to alanine in the crystallized structure. (D) Open-book representation of the RBD-TMPRSS2^S441A^ complex. The contact surface was colored indicating residues that form polar interactions (dark blue for RBD residues, light magenta for TMPRSS2), hydrophobic and aromatic residues (white), and residues that are at the interface and establish van der Waals contacts (light blue for the RBD and pink for TMPRSS2^S441A^). (E) Scheme highlighting the residues buried at the RBD-TMPRSS2^S441A^ interface (blue/pink shade). Amino acids that form salt bridges are indicated with filled triangles, while those that form hydrogen bonds are indicated with the empty symbol. For comparison, the sequence from the HKU1A RBD (isolate N1) is aligned below the one from HKU1B. Residues that are important for TMPRSS2 activity are indicated with stars (purple: catalytic triad; red: S1 site; orange: S2 site). Loops and β-strands are indicated, as in the previous panels. The numbers above the sequences correspond to the position in each polypeptide. (F) Comparison between the structures of different coronavirus RBDs (light blue) in complex with their receptors (dark pink). Two examples of betacoronaviruses (SARS-CoV-2 in complex with ACE2, PDB: 6M0J; MERS-CoV in complex with DPP4, PDB: 4L72), two of alphacoronaviruses (NL63-CoV with ACE2, PDB: 3KBH; HCoV-229E with APN, PBD: 6U7G) and one of a deltacoronavirus (PDCoV and human APN, PDB: 7VPQ) were selected. The buried surface area (BSA) of each complex is indicated below. (G) The structure of the RBD+TMPRSS2^S441A^ complex was aligned to the RBD from the closed HKU1A Spike (apo-closed, PDB: 8OHN), the closed Spike with a disialoglycan (Sia) in the NTD (holo-closed, PDB: 8OPM) and the Spike in an open form with a disialoglycan (Sia) (holo-1-RBD-up, PDB: 8OPN). An inset on the first panel zooms into the region indicated with an oval to better visualize the clashes between TMPRSS2 bound to a protomer in the ‘down’ conformation and amino acids and glycans from the other chains. Each protomer of the Spike is indicated with different colors (dark blue, light blue and white) and glycans are colored in green. (H) 293T cells expressing either GFP1-10 or GFP11 (GFP split system) were transfected with TMPRSS2 and HKU1B (either wild-type or harboring the W89A mutation). Cell-cell fusion was quantified by measuring the GFP area after 20 h. Data are mean ± s.d. of three different experiments. The dotted line indicates a normalized fusion of 1.0 relative to wild-type Spike. Statistical analysis: (H) One-way ANOVA with Dunnett’s multiple comparison test compared to the WT TMPRSS2 on non-normalized log-transformed data. *** p<0.001.

**Table 1.** Data collection and refinement statistics.

| PDB code | RBD+TMPRSS2+VHH-A01<br>XXX3 | TMPRSS2+VHH-A07<br>XXX1 | TMPRSS2+VHH-A07<br>XXX2 |
| --- | --- | --- | --- |
| **Data Collection** |  |  | (staraniso) |
| Space Group | P 6<sub>1</sub> | C 1 2 1 | C 1 2 1 |
| a, b, c (Å) | 201.9, 201.9, 210.3 | 182.7, 53.8, 65.5 | 161.9, 54.5, 165.8 |
| a, b, g (deg) | 90.0, 90.0, 120.0 | 90.0, 100.7, 90.0 | 90.0, 108.4, 90.0 |
| Resolution (Å) | 25-3.55 (3.64-3.55) | 25-1.80 (1.85-1.80) | 25-2.30 (2.39-2.30) |
| Rmerge | 0.265 (12.876) | 0.106 (2.643) | 0.240 (2.236) |
| Mean(I)/sd(I) | 14.6 (0.4) | 9.2 (0.6) | 6.6 (0.9) |
| Number of reflections | 58457 (4358) | 58045 (4194) | 50880 (2545) |
| Completeness (%) | 99.6 (100.0) | 99.8 (98.3) | 82.7 (39.4) |
| Comp. (ellipsoidal) |  |  | 95.0 (79.2) |
| Multiplicity | 43.3 (41.4) | 6.4 (6.3) | 7.0 (6.9) |
| CC(1/2) | 1.00 (0.223) | 0.998 (0.293) | 0.990 (0.316) |
| **Refinement** |  |  |  |
| Resolution (Å) | 25 - 3.55 | 25 - 1.80 | 25 - 2.30 |
| Number of reflections | 58058 | 57982 | 50824 |
| Rwork / Rfree | 19.29 / 22.13 | 17.69 / 21.54 | 20.87 / 24.71 |
| NCS | 2 |  | 2 |
| No. of atoms |  |  |  |
| Protein | 12567 | 4003 | 7032 |
| Ligand / Carb / Water | 95 | 389 | 207 |
| B-factor |  |  |  |
| Clashscore | 6.67 | 3.70 | 3.92 |
| R.m.s deviations |  |  |  |
| Bond lengths (Å) | 0.004 | 0.010 | 0.002 |
| Bond angles (°) | 0.788 | 1.013 | 0.493 |
| Ramachandran |  |  |  |
| Favored (%) | 96.05 | 96.15 | 96.52 |
| Outliers (%) | 0.13 | 0.20 | 0.11 |
| Rotamer outliers | 1.00 | 0.46 | 0.26 |

Diffraction limit #1: 2.796:( 0.8883, 0.0000, 0.4594): 0.984 **a\*** + 0.177 **c\***
Diffraction limit #2: 2.101:( 0.0000, 1.0000, 0.0000): **b\***
Diffraction limit #3: 2.241:(-0.4594, 0.0000, 0.8883):-0.413 **a\*** + 0.911 **c\***

**Table 2.**
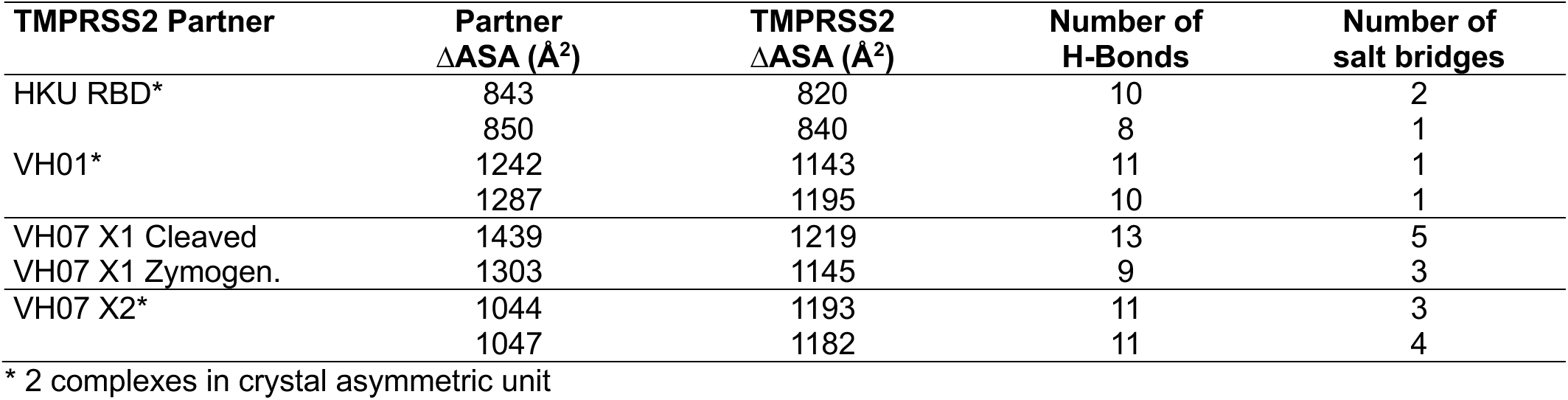
Buried surface area and number of polar interactions in the crystal structures.

| **TMPRSS2 Partner** | **Partner**<br>**ΔASA ($\text{\AA}^2$)** | **TMPRSS2**<br>**ΔASA ($\text{\AA}^2$)** | **Number of**<br>**H-Bonds** | **Number of**<br>**salt bridges** |
| --- | --- | --- | --- | --- |
| HKU RBD* | 843 | 820 | 10 | 2 |
|  | 850 | 840 | 8 | 1 |
| VH01* | 1242 | 1143 | 11 | 1 |
|  | 1287 | 1195 | 10 | 1 |
| VH07 X1 Cleaved | 1439 | 1219 | 13 | 5 |
| VH07 X1 Zymogen. | 1303 | 1145 | 9 | 3 |
| VH07 X2* | 1044 | 1193 | 11 | 3 |
|  | 1047 | 1182 | 11 | 4 |

\* 2 complexes in crystal asymmetric unit

### Highly complementary interacting surfaces

The HKU1B RBD/TMPRSS2 interface buries a surface area of about 1600 Å^2^ (∼800 Å^2^ on each side, Table 2) and has a shape complementarity (Sc) value of 0.72, which is substantially higher than the typical 0.64–0.68 range observed for antibody–antigen complexes ^27^. The polar network at the interface involves salt bridges and several hydrogen bonds (listed in Table SI1 and displayed in Figure 1C, left panel). The polar interactions surround a central hydrophobic patch in the surfaces of both RBD and TMPRSS2 (Figure 1D) which gives rise to a non-polar cluster at center of the interface (Figure 1C, right panel). We refer from hereon to RBD or TMPRSS2 residues using suffix R or T, respectively. Of note, residues W515_R_ and R517_R_, which, as we showed previously ^1^, abolish the interaction with TMPRSS2, are located at the interface. The residues of the catalytic triad are not directly contacted by the RBD, which instead reaches residues of the substrate binding sites (K342_T_ from the S1 site and T341_T_ at S2, in loop C), or that are very close to them (i.e. N433_T_ and S463_T_ with respect to D435_T_ and S436_T_ and G464_T_ in the S1 site; Figure 1E). Comparison with the mode of binding of the RBD of other coronaviruses to their receptor (Figure 1F) showed that the HKU1B RBD shares a similar buried surface area at the interface, although in the other coronaviruses the RBDs lack the pincer plier structure observed in HKU1.

### Only an RBD in the “up” conformation in an open spike can bind TMPRSS2

To understand the interactions of the HKU1B RBD with TMPRSS2 in the context of the trimeric spike, we superimposed the X-ray structure of the complex on the RBD of the HKU1-CoV spike ^12,28^. This exercise showed that TMPRSS2 cannot bind the RBD in the closed conformation because of clashes with the adjacent S1 protomers in the trimer (Figure 1G). In contrast, TMPRSS2 binding is unencumbered with the RBD in the “up” conformation, which is achieved only upon binding a disialoside glycan (9-O-Ac-Sia(α2,8)-Sia) in an allosteric pocket present in the N-terminal domain (NTD) of S1 ^12^. Our RBD/TMPRSS2 structure therefore predicts that mutations in the allosteric pocket that abolish binding of sialo glycans would result in a spike protein unable to bind TMPRSS2. We tested this prediction by using a split-GFP assay for syncytia formation in TMPRSS2-expressing cells ^1^. We introduced the W89A mutation in the NTD, which prevents sialo glycan binding ^10^ and generates spike proteins that display the same closed conformation as the wild-type spike in the apo form, as determined by cryo-EM ^12^. In line with these observations, the HKU1B W89A spike protein did not induce syncytia, despite normal surface expression (Figure 1H and Figure SI2). These data confirmed that TMPRSS2 acts as entry receptor only after the allosteric conformational change induced on the spike by sialo glycan binding to the NTD.

### Functional analysis of the RBD-TMPRSS2 interface

#### Mutagenesis of the HKU1-CoV spike protein

To establish the importance of the residues at the interface of the HKU1B RBD/TMPRSS2 complex, we used a battery of assays (Figure 2). We introduced mutations in plasmids coding for the HKU1B spike, generating 16 mutants that were properly expressed in 293T cells as assessed by flow cytometry (Figure SI2A). We co-transfected the spike mutants with TMPRSS2 to evaluate their fusogenic properties in the syncytia formation assay. Most of the tested spike variants showed a significantly reduced ability to induce cell-cell fusion, especially when the mutations involved residues that form salt bridges (K487_R_ and D505_R_) or are part of the hydrophobic cluster formed at the interface (L510_R_, V512_R_, L521_R_, Y528_R_) (Figure 2A). The effects on cell-cell fusion were further confirmed by introducing selected mutations on soluble RBDs and testing the binding to TMPRSS2^S441A^ by biolayer interferometry (BLI) (Figure 2B). Overall, the mutants showed diminished binding to the receptor measured at the stationary state, with L510R_R_, Y528A_R_ and K487A_R_ having the most significant effects.

**Figure 2.**
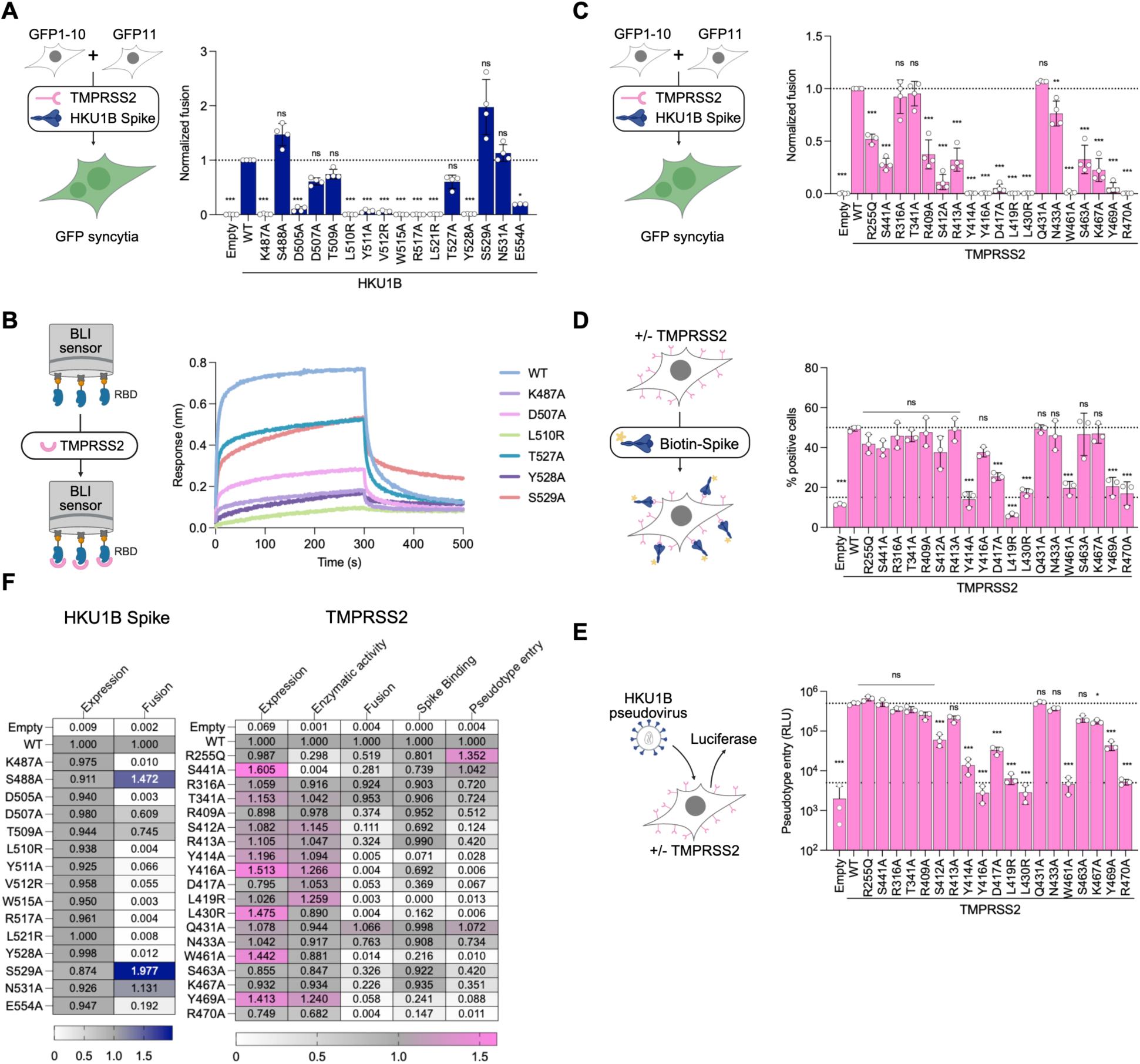
Functional experiments validate the RBD-TMPRSS2 crystal structure. (A) 293T cells expressing either GFP1-10 or GFP11 (GFP split system) were transfected with HKU1B Spike and TMPRSS2, and fusion was quantified by measuring the GFP area after 20 h. Data are mean ± s.d. of three different experiments. The dotted line indicates a normalized fusion of 1.0 relative to wild-type TMPRSS2. (B) Biolayer interferometry (BLI) experiment performed with soluble HKU1B RBDs containing mutations in the binding interface. The RBDs were immobilized to Ni-NTA sensors and their interaction with soluble TMPRSS2^S441A^ (800 nM) was followed in real-time. One representative experiment of three is presented. (C) Cell-cell fusion experiment carried out transfecting the wild-type HKU1B Spike with TMPRSS2 WT or different mutants. Fusion was quantified by measuring the GFP area after 20 h. Data are mean ± s.d. of four different experiments. The dotted line indicates a normalized fusion of 1.0 relative to wild-type TMPRSS2. (D) 293T cells were transfected with plasmids coding for TMPRSS2 or mutated variants, they were incubated with biotinylated soluble HKU1B Spike and binding to cells was determined by flow cytometry using labeled streptactin. Data are mean ± s.d. of three different experiments. The upper dotted line indicates the percentage of transfected cells with wild-type TMPRSS2 that were considered positive, while the bottom dotted line shows the background levels in non-transfected cells. (E) 293T cells transfected with TMPRSS2 WT or mutant variants were infected by luciferase-encoding HKU1B pseudoviruses. Luminiscence was read 48 h postinfection. RLU, relative light unit. The upper dotted line indicates the mean RLU obtained when cells were transfected with wild-type TMPRSS2, while the lower dotted line shows the background levels in non-transfected cells. (F) Heatmaps summarizing the functional data from panels A to E, along with the results from controls of expression and enzymatic activity (supplementary information) Statistical analysis: (A, C, E) One-way ANOVA with Dunnett’s multiple comparison test compared to the WT on non-normalized log-transformed data. (D) One-way ANOVA with Dunnett’s multiple comparison test compared to the WT. * p<0.05, ** p<0.01, *** p<0.001.

#### Design and analysis of TMPRSS2 mutants

We assessed the effects of TMPRSS2 mutations within the identified HKU1B interface on syncytia formation (Figure 2C), binding of soluble HKU1B spike (Figure 2D) and HKU1B pseudovirus entry (Figure 2E). We generated 19 mutants that had similar or even slightly higher cell surface expression levels than the wild-type protein, as determined by flow cytometry (Figure SI2B). Because the mutations could affect the protease activity required to cleave the spike, we tested each TMPRSS2 variant by measuring its ability to trigger fusion of the H229E-CoV spike after binding to its receptor, aminopeptidase N (APN) ^29^. Co-expression of 229E spike protein with APN generated syncytia in the presence of wild-type TMPRSS2 and not with the catalytically inactive TMPRSS2^S441A^ mutant (Figure SI2B), indicating that 229E spike cleavage by TMPRSS2 is required for fusion. This reporter system indicated that the protease activity of the designed TMPRSS2 mutants was not significantly affected. Co-transfecting the wild type HKU1B spike with TMPRSS2 variants showed that essentially all mutations at the receptor binding interface had a significantly reduced capacity to induce HKU1B-mediated cell-cell fusion (Figure 2C). As expected, the control mutant R316A_T_, located in loop E and not involved in receptor binding, induced normal cell-cell fusion levels.

We also analyzed by flow cytometry the binding of soluble HKU1B spikes to 293T cells expressing the TMPRSS2 variants. Most of the mutants impaired in cell-cell fusion showed decreased binding (Figure 2D). The most affected mutants were those that abrogated salt-bridges (D417A_T_ and R470A_T_) or hydrogen bonds (Y469A_T_) or disrupted the hydrophobic cluster at the interface (L410R_T_, Y414A_T_, L430R_T_, W461A_T_). The control R316A_T_ showed normal binding levels. We also used TMPRSS2-expressing cells to determine their sensitivity to lentiviral pseudotypes bearing the HKU1B spike (Figure 2E). We previously reported that mutations abrogating TMPRSS2 protease activity (R225Q and S441A) mediate viral infection at similar levels as the wild-type protein because the pseudoviruses can follow an endocytic route of entry ^1^. While the mutation R316A_T_ did not affect pseudovirus infection, the TMPRSS2 variants that showed reduced binding to the soluble spike were also inefficient for viral entry. The Y416A_T_ mutant did not trigger pseudovirus entry (Fig 2E), nor induced cell-cell fusion (Fig 2C).

Overall, as summarized in the heatmap of Figure 2F, our characterization of about 40 point mutants indicates that changes in residues at the interface of the HKU1B-RBD/TMPRSS2 complex impair binding, fusogenicity and infection, validating the crystal structure.

### TMPRSS2 residues at position 417 and 469 are key to define host tropism

We identified six positions in the TMPRSS2 SP domain -Y414_T_, Y416_T_, D417_T_, L419_T_, L430_T_, W461_T_, L469_T_ and R470_T_ - as the most relevant for the interaction with the HKU1B RBD. We then examined the pattern of conservation of these residues across different TMPRSS2 orthologs by aligning 201 mammalian TMPRRS2 sequences between amino acids 340 and 473 (numbering corresponding to human TMPRSS2), which span the interface with the HKU1B RBD. The resulting sequence logo (Figure 3A) indicated that among the residues listed above, Y416_T_, L419_T_, L430_T_ and W461_T_ are strictly conserved, while Y414_T_ is more than 80% conserved. In contrast, the variability is high at positions 417 and 469, where D and Y, the respective residues in humans, are not the most frequently found. Many species have a non-charged residue replacing D417, while polar (N) or non-polar (L, F) residues replace Y469. These observations suggest that residues at positions 417 and 469 may be determinants of TMPRSS2 function as HKU1 receptor in different mammalian species. We tested this hypothesis by assessing the capacity of TMPRSS2 from selected mammals (macaque, mouse, hamster, and ferret), to induce HKU1-CoV cell-cell fusion and pseudovirus infection. The respective TMPRSS2s display [D;N], [N;L], [N;L], and [N;N] instead of [D;Y] as in humans at positions [417;469] (Figure 3D). Despite similar expression levels (Figure SI2C), only human TMPRSS2 induced syncytia in 293T cells co-expressing the HKU1B spike (Figure 3B, left panel). The macaque TMPRSS2 (which differs only at position 469), allowed reduced pseudovirus entry (Figure 3B, right panel). Introducing the N469Y mutation into macaque TMPRSS2 restored syncytia formation and increased pseudovirus infection by ∼10-fold (Figure 3C). Changing Q467 on the macaque protein to the mostly conserved lysine did not show further significant effects on cell-cell fusion or pseudotype entry (Figure 3C). With mouse TMPRSS2, only the double mutant engineered to change the [N;L] to the human [D;Y] motif restored syncytia formation and pseudovirus entry, while the two single mutants were poorly or not functional (Figure 3C).

**Figure 3.**
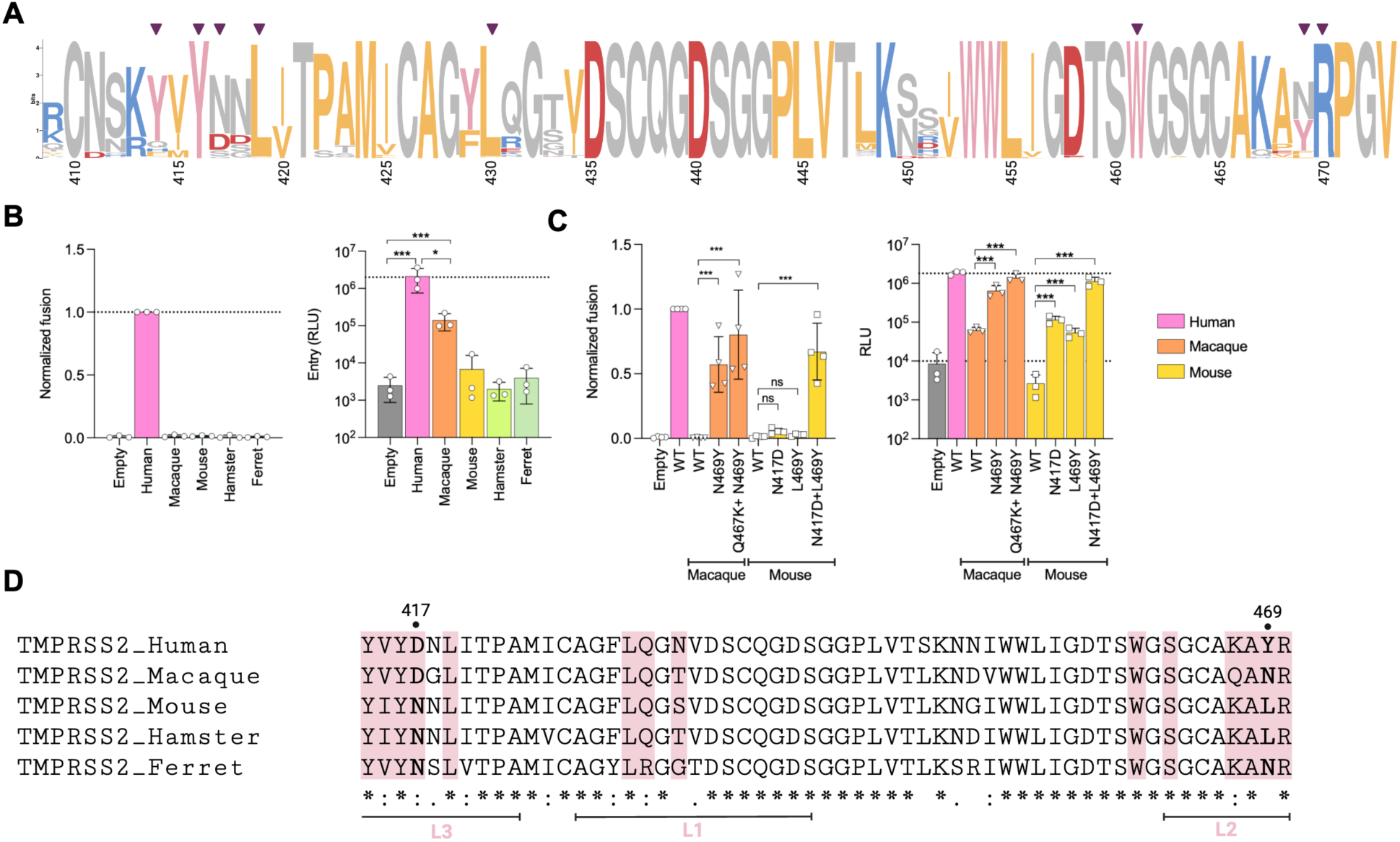
TMPRSS2 residues at positions 417 and 469 influence binding to different homologues. (A) Frequency of different amino acids occupying positions 409 to 473 (numbers at the bottom) of TMPRSS2 orthologs. The logo was obtained using 201 sequences from mammals. Purple inverted triangles indicate the positions identified as relevant for human TMPRSS2 in functional experiments (Figure 2). (B) Cell-cell fusion (left panel) and pseudovirus entry (right panel) experiments performed with cells transfected with TMPRSS2 from different species. Data are mean ± s.d. of three different assays. The dotted line indicates a normalized fusion of 1.0 relative to human TMPRSS2 or the mean RLU obtained with human TMPRSS2. (C) Cell-cell fusion (left panel) and pseudovirus entry (right panel) experiments performed with cells transfected with wild-type or mutant TMPRSS2 from selected mammals. Data are mean ± s.d. of three different experiments. The dotted line indicates a normalized fusion of 1.0 relative to human TMPRSS2 or the mean RLU obtained with human TMPRSS2. (D) Sequence alignment of the TMPRSS2 homologues used for the experiments in panel (B), highlighting the residues that are part of the interface formed with the HKU1-RBD (pink shade) and indicating TMPRSS2 loops L1 (loop 1), L2 (loops 2), L3 (loop 3) and LC (loop C). Statistical analysis: (B, C) One-way ANOVA with Šídák’s multiple comparison test * p<0.05, *** p<0.001.

### Nanobody A07 inserts its CDR3 into the TMPRSS2 substrate-binding groove

We next crystallized the complex between TMPRSS2^S441A^ and the inhibitory nanobody A07 (Figure 4A, Figure SI3) and determined its structure to 1.8 Å resolution (Table SI1). A07 covers the active site cleft and buries a large surface area (about 2600 Å^2^, ∼1400 Å^2^ on the VHH and ∼1200 Å^2^ on TMPRSS2), interacting with residues in exposed loops of the SP domain (loops 1, 2, 3, A, B, C, D), some also involved in the interaction with HKU1B RBD (Figure 4A). Superimposing the RBD+TMPRSS2^S441A^ structure on the A07+TMPRSS2^S441A^ complex showed clashes between the nanobody and the RBD (Figure 4B), explaining the blocking activity and providing further validation to the interaction site that we report here for the RBD. Nanobody A07 contacts TMPRSS2 almost exclusively through its CDRs. The most notorious feature is the insertion of its long (21 residues) CDR3 in the substrate binding cleft in between the two lobes of the SP domain. Superposing the complex with the peptide-bound structures of hepsin and TMPRSS13 shows that the side chain of R103_A07_ occupies the P1 position, making contacts with residues D435_T_, S436_T_ and G464_T_ (Figure 4C), which form the S1 site ^18^.

**Figure 4.**
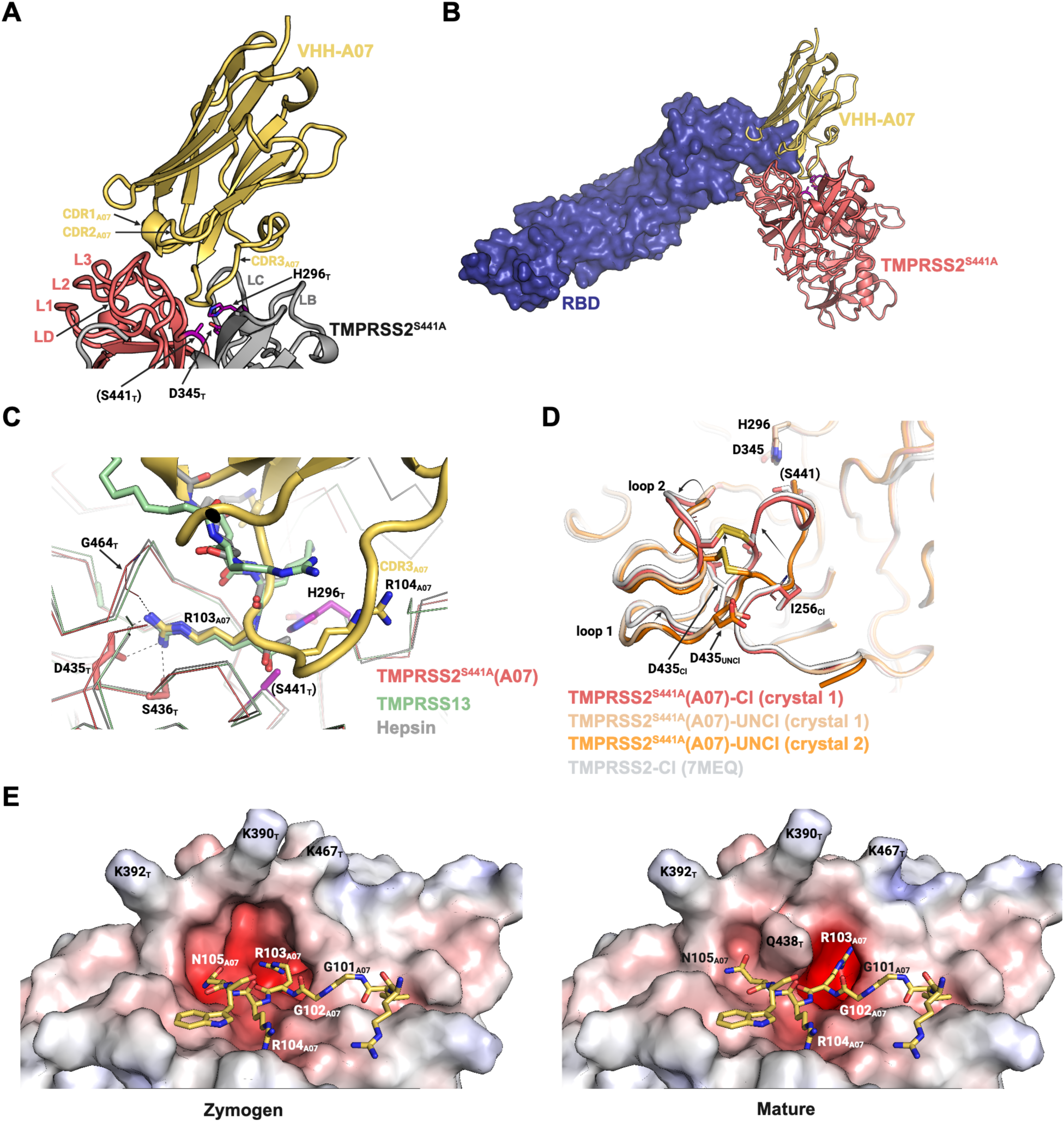
VHH-A07 blocks TMPRSS2 activity and binding to the HKU1-RBD by inserting its CDR3 in the active site. (A) Crystal structure of the nanobody VHH-A07 (yellow) complexed to TMPRSS2^S441A^. The two β-barrels that form the SP domain are colored in gray (β-barrel 1) and light red (β-barrel 2). Important structural elements are indicated in the nanobody (CDR1_A07_, CDR2_A07_, CDR3_A07_), as well as in TMPRSS2^S441A^ (loops B, C, D, 1, 2, 3). Residues from the catalytic triad are shown in purple. Subscripts in the labels identify the protein. The active site S441 was mutated to alanine in the crystallized construct, so it is annotated between parentheses. (B) Superposition of the TMPRSS2^S441A^+RBD and TMPRSS2^S441A^+VHH-A07 complexes. For simplicity, TMPRSS2 from the complex with the RBD is not shown. (C) Superposition of TMPRSS2^S441A^ in complex with VHH-A07 (protease in light red, nanobody in yellow) with the structures of TMPRSS13 (green, PDB: 6KD5) and Hepsin (gray, PDB: 1Z8G) bound to substrate-analog inhibitors (decanoyl-RVKR-chloromethylketone shown in green sticks and Ac-KQLR-chloromethylketone in gray ticks, respectively). For clarity, the proteases are shown in ribbon representation and the nanobody in cartoon. Residues from the TMPRSS2 catalytic triad are indicated in purple, with S441 labeled with parenthesis to indicate that it is mutated to alanine in the crystalized construct. Residues that form the S1 site (D435, S436, G464) are indicated, as well as R103 from nanobody A07. (D) Superposition of different TMPRSS2^S441A^ structures obtained in complex with VHH-A07. A first crystal form (crystal 1) allowed building two atomic models, where one has the characteristic features of a cleaved (Cl, light red) serine protease, while the other represents the zymogen (UNCl, wheat) form. The second crystal form (crystal 2) allowed building the model shown in orange that corresponds to uncleaved TMPRSS2^S441A^. A previously reported structure of active TMPRSS2 is shown in white for comparison (PDB: 7MEQ). The I256 residue, which follows the autocleavage site, is indicated. Important TMPRSS2 loops are designated as well as the positions of the catalytic triad (H296, D345, S441A) and D435 from the S1 site. The disulfide C437-C465 is shown in yellow sticks. Arrows with tapered lines indicate the repositioning on relevant segments upon cleavage. (E) Comparison between the TMPRSS2^S441A^ active site in the uncleaved (left) and cleaved (right) forms found in crystal 1. The surface of the protease was colored according to the electrostatic potential and the residues R99_A07_-W106_A07_ from A07 CDR3 are shown in sticks.

The TMPRSS2^S441A^/A07 crystals displayed electron density for the TMPRSS2 LDLR-A domain, which was not resolved in the previous structure of TMPRSS2 ^18^. The LDLR-A domain includes a calcium-binding site formed by the side chains of D134_T_, H138_T_, D144_T_, and E145_T_, and the main chain carbonyl group of N131_T_ and V136_T_ (Figure SI4A).

The SP domain of cleaved TMPRSS2 (TMPRSS2-Cl) crystallized previously in complex with the Nafamostat inhibitor ^18^, superposes very well with its counterpart in the TMPRSS2^S441A^/A07 crystals, with a root mean square deviation (RMSD) of 0.315 Å for 1502 atoms in 236 residues. In this structure, the residues immediately downstream the autocleavage site (I256_T_ and V257_T_) are found in an internal pocket where the free amino group of I256_T_ forms a salt-bridge with the side chain of D440_T_ (Figure 4D). This interaction can only be established after cleavage of the protease and is a characteristic feature of serine proteases in the active conformation ^30^. This observation indicates that TMPRSS2^S441A^ in the crystals of the complex with A07 underwent cleavage by a contaminating protease. The crystallographic data further indicated that the maturation cleavage did not occur in 100% of the molecules forming the crystal. Another conformation was detected, for which the high-resolution diffraction allowed the refinement of an atomic model (Table 1). In this second model, the loop bearing the cleavage site was disordered, with the first residue with clear electron density after the cleavage site being G259_T_, as expected for an uncleaved form – or zymogen - of TMPRSS2^S441A^. We confirmed that this is indeed the case by crystallizing the TMPRSS2^S441A^/A07 complex under different conditions, which yielded crystals that diffracted to 2.4 Å resolution. The residues I256_T_ and V257_T_ were not visible in the structure determined from these new crystals, in which TMPRSS2^S441A^ showed no evidence of proteolysis. The resulting model aligned very well with the second conformation described above, supporting the hypothesis that it corresponds to the TMPRSS2 zymogen. Furthermore, this structure has the characteristic “zymogen triad” initially observed for chymotrypsinogen, D194-H40-S32 ^31^, corresponding to D440-H279-S272 in TMPRSS2 (Figure SI4B). Upon activation, D440 is released from this triad to make the salt-bridge/hydrogen bond with the newly formed N-terminus at I256. This rearrangement leads to formation of the oxyanion hole required for cleavage of the scissile peptide bond of the substrate. Proteases that do not have a zymogen triad, such as the tissue-type plasminogen activator, have a high level of catalytic activity in the zymogen form ^32^.

Comparison of the structures of cleaved and zymogen TMPRSS2^S441A^ reveals that the main changes are localized to loops 1 (L430_T_-D440_T_) and 2 (G462_T_-K467_T_). In the zymogen, residues 430-440 occupy an external position and S1 site residues (D435_T_, S436_T_) are away from the active site (Figure 4D).

As mentioned above, in TMPRSS2^S441A^-Cl the free N-terminus of I256_T_ flips to the interior of the molecule, displacing the 430-440 segment towards the core of the domain (Figure 4D). The disulfide bond between C437_T_ and C465_T_ propagates this movement to loop 2 (G462_T_-K467_T_), which also adopts a different conformation (Figure 4D). As a consequence, the S1 site is formed without alteration of the catalytic triad. Of note, the conformation of loops 1 and 2 in TMPRSS2^S441A^-Cl is the same those of active forms of hepsin ^33^ and TMPRSS13 (also called MSPL) ^34^ (Figure SI4C).

### TMPRSS2 maturation affects binding affinity towards the HKU1-CoV RBD

As described above, maturation of TMPRSS2 into an active form implies changes in the conformation of loops 1 and 2. Since they are part of the HKU1B RBD binding site, we hypothesized that TMPRSS2 cleavage may impact the interaction with the RBD. We therefore conducted BLI experiments with immobilized RBD, soluble TMPRSS2^S441A^ and cleaved soluble TMPRSS2^S441A^ (TMPRSS2^S441A^-Cl). The latter was obtained by previous digestion of the zymogen with the wild-type protease. SDS-PAGE confirmed that the digestion of TMPRSS2^S441A^ was complete, generating fragments of ∼25-30 kDa associated by a disulfide bond (Figure SI5). The BLI curves showed that TMPRSS2^S441A^-Cl displays a dramatic increase in the association signal and reduced dissociation rate (Figure 5A). The curves corresponding to TMPRSS2^S441A^-Cl fitted well to a 1:1 binding model. We determined the kinetic association and dissociation rates, and measured a Kd of 30 nM (Figure 5C). This value is ̃5-fold higher than the one obtained for TMPRSS2^S441A^-zymogen ^1^. We also used TMPRSS2^S441A^-Cl to determine by BLI the affinity constants of some of the RBD mutants (Figure 5B). This confirmed that K487A_R_, L510R_R_, and Y528A_R_ were the most impaired (Figure 5C). The mutant S529A_R_ showed increased affinity, mostly due to a lower dissociation rate.

**Figure 5.**
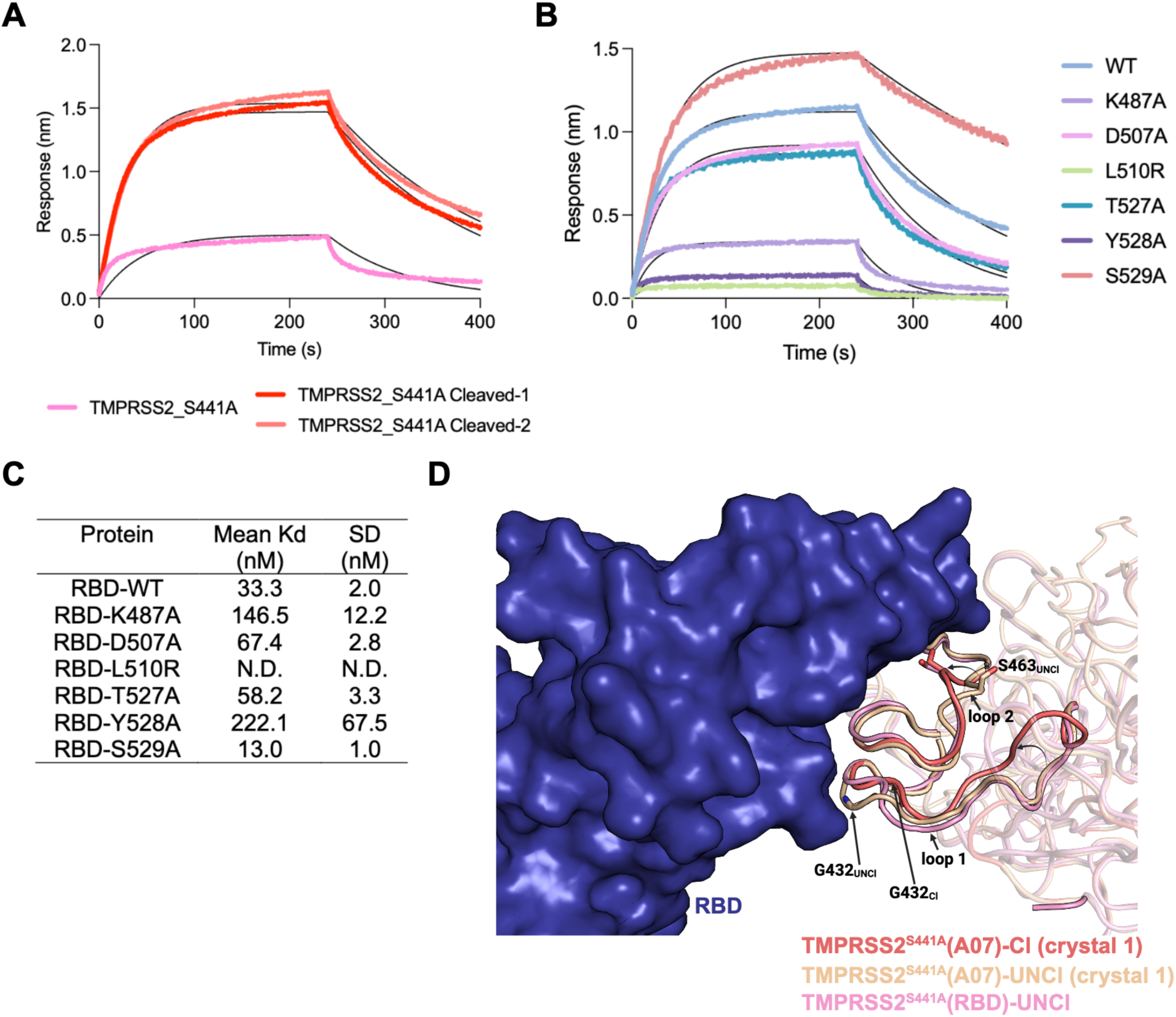
TMPRSS2 cleavage affects binding to the HKU1 RBD. (A) Curves of a BLI experiments performed immobilizing the HKU1B RBD and measuring the response upon incubation with 150 nM TMPRRS2^S441A^ or with 150 nM TMPRSS2^S441A^ previously cleaved by incubation with the wild-type protease. Two separate cleavage reactions were carried out. The colored curves correspond to the experimental data and the thin black lines are the curves fitted to a 1:1 binding model. One representative experiment of three is presented. (B) BLI curves of experiments performed immobilizing different RBDs and measuring the response after incubation with 120 nM cleaved TMPRSS2^S441A^. The colored curves correspond to the experimental data and the thin black lines are the curves fitted to a 1:1 binding model. One representative experiment of three is presented. (C) Kd values of different RBDs towards cleaved TMPRSS2^S441A^. Three experiments were performed for each RBD using a TMPRSS2 concentration range from 7.5 to 240 nM. (F) Superposition of the TMPRSS2^S441A^ structure determined in complex with the HKU1-RBD (TMPRSS2 in pink, RBD in blue surface) or in complex with VHH-07 (crystal 1: atomic model of the cleaved form shown in light red, and the zymogen form in wheat). Loops that form the binding site and change upon cleavage (loops 1 and 2) are labeled. The positions where conformational changes were observed are indicated with the residues that are implicated (backbone atoms of G432 and side chain of S463). Arrows with tapered lines indicate the movement of relevant segments upon TMPRSS2 autocleavage.

We then compared the structures of TMPRSS2^S441A^-Cl and TMPRSS2^S441A^-zymogen (obtained with A07) with that of TMPRSS2^S441A^ in complex with the RBD (Figure 5D). In the latter structure, residues I256_T_ and V257_T_ were not visible, indicating that TMPRSS2^S441A^ was uncleaved. Overall, the protease conformation matched TMPRSS2^S441A^-zymogen, except for local differences in the loops at residues around G432_T_ and S463_T_. In the complex with the RBD, these loops adopt a conformation closer to the one found in TMPRSS2 ^S441A^-Cl, where G432_T_ moves towards the active site, further away from the RBD interface, and S463_T_ flips outwards to form a hydrogen bond with the RBD (Figure 5D) explaining why the affinity of TMPRSS2^S441A^- Cl for the RBD is higher than that of the zymogen.

## DISCUSSION

We report the structure of HKU1B RBD bound to its receptor TMPRSS2, revealing a unique binding mode among coronaviruses. The HKU1-CoV RBD uses its pincer plier motif at its distal end to recognize the TMPRSS2 substrate specificity loops. The interaction blocks access of substrates to the catalytic groove, inhibiting the proteolytic activity. This inhibition is further supported by the clashes observed between the HKU1B RBD and a nanobody that mimics an incoming substrate. Our results are in line with a recent report on the structure of HKU1A RBD complexed with cleaved TMPRSS2 and the inhibition of TMPRSS2 proteolytic activity by the HKU1A RBD ^35^

The HKU1-CoV spike protein binds cell-surface disialoside glycans (9-O-Ac-Sia(α2,8)Sia) through a binding pocket in the NTD, causing the allosteric opening of the spike ^12^. This led to the speculation that the sialo glycan is a primary receptor, while a proteinaceous receptor would be secondary. This mechanism ensures that the spike will not undergo premature activation before reaching a cell, limiting exposure of the receptor binding motif of the RBD to neutralizing antibodies. Moreover, when interacting with an up-RBD, the active site of TMPRSS2 is far from the spike core, indicating that a single TMPRSS2 molecule cannot simultaneously cleave the spike and act as receptor. By binding to up-RBD conformations, TMPRSS2 traps partially open intermediates of the HKU1 spike, displacing the conformational equilibrium towards the open form, where S2 is unshielded and primed for fusion, acting by the same mechanism as ACE2 with the SARS-CoV-2 spike ^36^. Considering that ∼45% of HKU1-CoV spike with (9-O-Ac-Sia(α2,8)Sia) were found closed or with only one RBD ‘up’ ^12^, a secondary protein receptor with high affinity towards the RBD would guarantee the unidirectionality of the conformational change towards the open spike.

We also determined the crystal structure of TMPRSS2 bound to nanobody A07, which competes with the RBD for binding ^1^. A07 inserts its long CDR3 in the active site cleft, with R103_A07_ occupying the P1 position (Figure 4E). It has been suggested that the predicted S1’ site (V280_T_, H296_T_, C297_T_) accepts small hydrophobic P1’ residues ^18^. Our structure shows R104_A07_ in the S1’ site and its side chain is well accommodated, interacting with main chain atoms from H296_T_ and C297_T_, without altering the conformation of the loop containing V280_T_. Given that the peptide bond R103_A07_-R104_A07_ lies in the right position to undergo the attack of the active site S441, it is possible that the nanobody is cleaved by TMPRSS2. Nevertheless, we expect the extensive network of polar interactions and the large BSA to sustain binding even if the antibody is cleaved.

The structures of cleaved and uncleaved forms of TMPRSS2 highlight the internal reorganization of the protease upon autocleavage. In the zymogen form, loop 1 adopts a conformation where S1 residues are away from the active site and not in an optimal position to allow substrate binding, as illustrated in Figure 4E (left panel). The proteolytic cleavage releases the N-terminus of I256_T_, which flips towards the interior of the protein, with formation of a salt bridge between the charged amino group of I256_T_ and D440_T_. In turn, loop 1 is displaced and adopts a new conformation, transmitting this movement to loop 2 via a disulfide bond connecting them. Consequently, the S1 site residues (D435_T_ and S436_T_ from loop 1, and G464_T_ from loop 2) are brought into place to allow substrate binding. These changes agree with the general activation mechanism proposed for other serine proteases ^37^. Furthermore, the TMPRSS2 loops of the cleaved form adopt the same position as other enzymes of the same family (hepsin and TMPRSS13), suggesting they adopt a stable conformation in the active form. The structure of the A07/TMPRSS2 zymogen further shows how the substrate mimicking CDR3 is accommodated in the groove in the absence of the S1 site (Figure 4E, right panel), which forms only after activation. Together with the inherent plasticity of the zymogen, this data can potentially inform the development of TMPRSS2-specific small molecules targeting this site.

We also established that TMPRSS2 activation affects interaction with the HKU1-CoV RBD. In zymogen TMPRSS2^S441A^, the residues 431-433 (loop 1) adopt a conformation that may not favor binding to the RBD, which has the j2 jaw of the pincer plier facing it. Moreover, S463 in loop 2 is too far to interact with D507_R_. Upon cleavage, the 431-433 segment moves towards the core of TMPRSS2, facilitating the approach of the RBD, and S463 moves outwards, forming a hydrogen bond with D507_R_. These changes in loops 1 and 2 appear to be important for binding, since the structure of zymogen TMPRSS2^S441A^ with the RBD shows that they adopt the position similar to that observed in activated TMPRSS2^S441A^-Cl. This ‘mixed’ conformation, with local changes restricted the segments 431-433 and 463-464 might represent a high-energy state, which would decrease the affinity. In addition, the zymogen TMPRSS2^S441A^ loops could adopt the ensemble of observed conformations, having a higher entropic content than TMPRSS2^S441A^-Cl and resulting in lower affinity towards the HKU1-CoV RBD. The BLI curves show that zymogen TMPRSS2^S441A^ does not fit to a 1:1 binding model, while TMPRSS2 ^S441A^-Cl does, in line with a heterogeneous population of conformations that becomes homogeneous upon cleavage.

Our study has limitations. Firstly, we did not study the structure of the interaction of HKU1A with TMPRSS2. We previously reported that both HKU1A and HKU1B bind to and use TMPRSS2 as a functional receptor^1^. It is thus likely that the two HKU1 spikes similarly interact with TMPRSS2. The recent description by cryoEM of a TMPRSS2/ HKU1A RBD complex ^35^, revealed a similar mode of interaction for both HKU1A and B. Secondly, we did not solve the structure of the whole spike in association with its receptor. We circumvented this problem by superposing the structure of the RBD in complex with TMPRSS2 to the known structure of the HKU1 spike in order to characterize the conformational changes observed in both the spike and its receptor during viral binding and fusion.

In sum, the structure of the HKU1B RBD in complex with TMPRSS2 provides insights into the mode of action of the receptor and on the mechanism of spike activation. We also identify the most important residues of the interface, highlighting how they contribute to host species tropism, paving the way to future characterization of potential animal reservoirs. Finally, we describe the maturation of TMPRSS2 from a zymogen to an active protease and how it impacts binding to HKU1-CoV. The observed plasticity of the TMPRSS2 zymogen stands out as a vulnerable feature of the protease that can be specifically targeted to prevent activation. Our results therefore open the way to the development of specific drugs against dysregulated TMPRSS2 in tumors without affecting other serine proteases to avoid toxicity.

## METHODS

### Construct design

#### For production of recombinant proteins

Codon-optimized synthetic genes coding for the HKU1B RBD (residues 330-614 of the spike protein, isolate N5P8, NCBI accession Q0ZME7), the RBD and subdomain-1 (RBD-SD1, residues 307-675) and human TMPRSS2 (residues 107-492, NCBI accession O15393) were obtained from Genscript. Cloning and mutagenesis of those genes were also performed by Genscript. The RBD was cloned into pCAGGS, following a murine immunoglobulin kappa signal peptide, and upstream of a thrombin cleavage site and *in-tandem* Hisx8, Strep and Avi-tags. RBD-SD1, TMPRSS2 wild-type (WT) and TMPRSS2^S441A^ were cloned into a modified pMT/BiP plasmid (Invitrogen; hereafter termed pT350), which translates the protein in frame with an enterokinase cleavage site and a double strep-tag at the C-terminal end. Selected VHHs have been previously obtained ^1^ and were cloned into the bacterial expression vector pET23 with a C-terminal His-tag and N-terminal Met-Ala residues introduced during subcloning.

#### For cell transfection and functional assays

Codon-optimized synthetic genes coding for the full-length spike of HKU1 B/C isolate N5P8 (referred to as HKU1B, UniProtKB/Swiss-Prot: Q0ZME7.1) and those coding for TMPRSS2 from mouse (Mouse C57BL/6 - UniPropt: Q3UKE3), ferret (Ferret – UniProt: A0A8U0SMZ2), hamster (Syrian Hamster Isoform X1 – UniProt: A0A1U8BWQ2) and macaque (Macaque – UniProt: F6SVR2) with an N-terminal cMYC-tag were ordered to GeneArt (Thermo Fisher Scientific) and cloned into a phCMV backbone (GeneBank: AJ318514) by replacing the VSV-G gene. pQCXIP-Empty control plasmid was previously described ^38^. pQCXIP-BSR-GFP11 and pQCXIP-GFP1-10 were a kind gift from Yutaka Hata ^39^ (Addgene plasmid #68716 and #68715). pCSDest-TMPRSS2 was a kind gift from Roger Reeves ^40^ (Addgene plasmid # 53887). Human pCCAGS N-terminal cMYC-epitope tagged TMPRSS2 was a kind gift from Stefan Pöhlmann ^41^. Mutations in the HKU1 spike and TMPRSS2 were introduced using the NEB Q5 Side-Directed mutagenesis kit. Plasmids were sequenced before usage.

### Cells

HEK293T (293T) were from ATCC and cultured in DMEM with 10% fetal bovine serum (FBS) and 1% penicillin/streptomycin (PS). GFP-split cells were previously described ^38^ and cultured with 1 µg/mL of puromycin (InvivoGen). Cells were routinely screened for mycoplasma. Cells were authenticated by genotyping (Eurofins).

### Protein expression and purification

#### Protein expression and purification for X-ray crystallography

Plasmids encoding RBD-SD1 or TMPRSS2^S441A^ were co-transfected with the pCoPuro plasmid for puromycin selection in Drosophila Schneider line 2 cells (S2). The cell lines underwent selection in serum-free insect cell medium (HyClone, GE Healthcare) containing 7 μg/ml puromycin and 1% penicillin/streptomycin. For protein production, the cells were grown in spinner flasks until the density reached 10^7^ cells/mL, at which point the protein expression was induced with 4 μM CdCl_2_. After 6 days, the cultures were centrifuged, and the supernatants were concentrated and used for affinity purification in a Streptactin column (IBA). The strep tags were removed by incubating the proteins with 64 units of Enterokinase light chain (BioLabs) in 10 mM Tris, 100 mM NaCl, 2 mM CaCl_2_, pH 8.0, at room temperature, overnight. The proteolysis reactions were buffer-exchanged into 10 mM Tris, 100 mM NaCl, pH 8.0, and subjected to a second affinity purification, recovering the flow-through fraction containing the untagged proteins. The proteins were concentrated and the enzymatic deglycosylation with EndoD and EndoH was set up at room-temperature following overnight incubation with 1000 units of each glycosidase in 50 mM Na-acetate, 200 mM NaCl, pH 5.5. The proteins were further purified on a size exclusion chromatography (SEC) Superdex 200 16/60 (Cytiva) column in 10 mM Tris, 100 mM NaCl, pH 8.0, and concentrated in VivaSpin concentrators.

E. coli BL21pLysS cells were transformed with the plasmids encoding the VHHs, which were expressed in the cytoplasm after overnight induction with 0.5 mM IPTG at 16° C. The cultures were centrifuged, the bacterial pellets were resuspended in 40 mL of lysis buffer (20 mM Tris-HCl, 200 mM NaCl, 20 mM imidazole, pH 8.0) containing complete protease inhibitor cocktail (Roche) and they were frozen at −80 °C until used. On the purification day, the resuspended pellets were thawed, sonicated (15 minutes, 9s on-pulse, 5s off-pulse), centrifuged and loaded onto a HisTrap column. Bound proteins were eluted with a linear gradient of buffer B (20 mM Tris-HCl, 200 mM NaCl, 500 mM imidazole, pH 8.0) and analyzed by SDS-PAGE. Fractions with higher purity were pooled, concentrated and further purified by SEC on a Superdex 75 16/60 column (Cytiva) pre-equilibrated in 10 mM Tris-HCl, 100 mM NaCl, pH 8.0.

The purity of the final protein samples was analyzed by SDS-PAGE followed by Coomasie Blue staining.

#### Purification of complexes used for crystallization screenings

The RBD-SD1 construct was incubated with TMPRSS2^S441A^ and A01 at final concentrations of 47.4 μM, 71.1 μM and 107 μM, respectively. After over-night incubation at 4 °C, the reaction was loaded onto a Superdex 200 10/300 column (Cytiva) equilibrated in 10 mM Tris-HCl, 100 mM NaCl (pH 8.0) to isolate the complex by SEC. Eluted fractions were analyzed by SDS-PAGE and those corresponding to the ternary complex were pooled, concentrated to 8.5 mg/mL and used in crystallization trials.

TMPRSS2^S441A^ was incubated with A07 at final concentrations of 68 μM and 102 μM, respectively, over-night at 4 °C. Then, the mix was loaded onto a Superdex 200 10/300 column (Cytiva) equilibrated in 10 mM Tris-HCl, 100 mM NaCl (pH 8.0) and eluted fractions were analyzed by SDS-PAGE. Fractions of the binary complex were pooled, concentrated to 6.1 mg/mL and used in crystallization trials.

#### Protein expression and purification for biophysical assays

RBD-encoding plasmids were transiently transfected into Expi293F^TM^ cells (Thermo-Fischer) using FectoPro^®^ DNA transfection reagent (PolyPlus). After 5 days at 37 °C, cells were harvested by centrifugation and proteins from the supernatants were purified using a HisTrap-Excel column (Cytiva). Eluted fractions were pooled, concentrated and injected onto a Superdex 200 10/300 column (Cytiva) equilibrated in 10 mM Tris-HCl, 100 mM NaCl (pH 8.0) to perform size-exclusion chromatography. Fractions from the main peak were concentrated and frozen.

TMPRSS2 (WT) was expressed from a stable S2 cell line as indicated before, and it was purified by affinity (the tag and glycans were not removed).

The purity of the final protein samples was analyzed by SDS-PAGE followed by Coomasie Blue staining.

#### Cleavage of TMPRSS2^S441A^ for biophysical assays

To prepare the first batch of cleaved TMPRSS2^S441A^, 600 µg were incubated at room temperature with 1 µg of TMPRSS2 WT in 600 µL of buffer (10 mM Tris-HCl, 100 mM NaCl, pH 8.0) for 21 hours. The final concentration of TMPRSS2^S441A^ in the reaction was 21 µM. Then, the mix was stored at 4 °C for 8 hours, aliquoted, flash-frozen in liquid nitrogen, and stored at −80 °C until used.

The second batch of cleaved TMPRSS2^S441A^ was prepared by incubating at room temperature 230 µg with 0.3 µg of TMPRSS2 WT in 200 µL of buffer (10 mM Tris-HCl, 100 mM NaCl, pH 8.0) for 7 hours. The final concentration of TMPRSS2^S441A^ in the reaction was 25 µM. Then, the mix was aliquoted, flash-frozen in liquid nitrogen, and stored at −80 °C until used.

### Crystallization and structural determination

The RBD-SD1+TMPRSS2^S441A^+A01 complex crystallized in 0.35 M NaH_2_PO_4_, 0.65 M K_2_HPO_4_ at 4°C using the sitting-drop vapor diffusion method. The TMPRSS2^S441A^+A07 complex crystallized in 20 %w/v PEG 3350, 0.05 M HEPES (pH 7.0), 1 %w/v Tryptone, 0.001 %w/v NaN3 (crystal form 1) and in 10 %w/v PEG 3000, 0.1 M imidazole (pH 8.0), 0.2 M lithium sulfate (crystal form 2) at 18 °C using the sitting-drop vapor diffusion method. Crystals were flash-frozen by immersion into a cryo-protectant containing the crystallization solution supplemented with 33% (v/v) glycerol, followed by rapid transfer into liquid nitrogen.

The X-ray diffraction data of both complexes were collected at the SOLEIL synchrotron source (Saint Aubin, France). Collections were carried out at 100 K at the Proxima-1 beamline ^42^.

Data were processed, scaled and reduced with XDS ^43,44^ and AIMLESS ^45^. The structures were determined by molecular replacement using Phaser from the PHENIX suite ^46^ with search ensembles obtained from AlphaFold2 (HKU1B-RBD-SD1 and A07) or from previously deposited structures (7MEQ for TMPRSS2^S441A^, 7KN5 for A01). The final models were built by combining real space model building in Coot ^47^ with reciprocal space refinement with phenix.refine.

The RBD/ TMPRSS2^S441A^/A01 complex crystallized in hexagonal P6_1_ space group and diffracted to 3.5 Å resolution with two ternary complexes in the asymmetric unit. Refinement was carried out using constraints provided by NCS and model targets (5KWB for the RBD, our high-resolution structure of TMPRSS2^S441A^ in complex with A07 for the TMPRSS2^S441A^ protein, and 7KN5 and 7VOA for the nanobody A01) allowing us to build a model with good geometry (Table 1).

The TMPRSS2^S441A^/A07 complex provided two crystal forms belonging to the C2 orthorhombic space group. One form diffracted to 1.8 Å with one complex in the asymmetric unit, and the other diffracted to 2.4 Å with 2 complexes in the asymmetric unit (Table 1). From the high-resolution crystal we built a model of cleaved TMPRSS2^S441A^, while the second crystal form allowed us to build the uncleaved form. Nevertheless, many strong positive peaks remained in the difference electron density map of the high-resolution data, particularly near the active site. These densities were easily explained by superposing the uncleaved TMPRSS2^S441A^ structure, clearly showing the presence of an alternative conformation on two loops in this crystal (residues 431-440 and 462-467, linked by the disulfide bond C437-C465), which refined with final occupancies of 0.55 and 0.45 (cleaved and uncleaved forms, respectively). In addition, the cleavage released a new N-terminus (residues 256-258) that became ordered and inserted deeply into the core of the protease to interact with the side chain of the buried D440, as it is observed in this family of proteases upon activation.

The final models were validated with Molprobity ^48^. The analyses of the macromolecular surfaces were carried out in PDBePISA ^49^. Figures were created using Pymol ^50^ and BioRender.com.

### Biolayer interferometry (BLI)

Affinity of recombinant RBDs towards the purified ectodomain of TMPRSS2^S441A^was assessed in real-time using a bio-layer interferometry Octet-R8 device (Sartorius). Nickel-NTA capture sensors (Sartorius) were loaded for 10 min at 1,000 rpm shaking speed with the different RBDs at 400 nM in PBS. The sensors were then blocked with PBS containing BSA at 1.0 mg/mL (assay buffer) and were incubated at 1,000 rpm with two-fold serially diluted concentrations (800 nM to 25 nM) of TMPRSS2^S441A^ in assay buffer. Association and dissociation were monitored for 300 s and 240 s, respectively. Measurements for a reference were recorded using a sensor loaded with an unrelated protein (CD147) that was dipped at each analyte concentration. A sample reference measurement was recorded from a sensor loaded with each RBD and dipped in the assay buffer. Specific signals were calculated by double referencing, substracting nonspecific signals obtained for the sensor and sample references from the signals recorded for the RBD-loaded sensors dipped in TMPRSS2^S441A^ solutions. Three independent experiments were performed but only the curves obtained at 800 nM in the first experiment were chosen for preparing figures.

Affinity of the recombinant RBDs towards cleaved TMPRSS2^S441A^ (first batch) was determined following a similar protocol, although the range of ligand concentrations assayed went from 240 nM to 7.5 nM. Association and dissociation were monitored for 240 s and 180 s, respectively. Specific signals were calculated by subtracting the nonspecific signal of the sample reference from the signals recorded for the RBD-loaded sensors. Association and dissociation profiles were fitted assuming a 1:1 binding model. Three independent experiments were performed and the Kd values from each of them were averaged and used to calculate the standard deviation.

### Sequence alignment

Multiple sequence alignments were performed using Clustal Omega ^51^. The sequence logo was created with WebLogo ^52^ (https://weblogo.berkeley.edu/).

### GFP-split fusion assay

Cell–cell fusion assays were performed as previously described ^38^. Briefly, 293T cells stably expressing GFP1-10 and GFP11 were co-cultured at a 1:1 ratio (6 × 10^4^ cells/well) and transfected in suspension with Lipofectamine 2000 (Thermo) in a 96-well plate (uClear, #655090) (20 ng of spike plasmid, 20ng of TMPRSS2 plasmids adjusted to 100 ng DNA with pQCXIP-Empty). At 20 h post-transfection, images covering 90% of the well surface, were acquired per well on an Opera Phenix High-Content Screening System (PerkinElmer). The GFP area was quantified on Harmony High-Content Imaging and Analysis Software.

### Pseudovirus generation and infection

Pseudoviruses were produced by transfection of 293T cells as previously described ^1^. Briefly, cells were cotransfected with plasmids encoding for lentiviral proteins, a luciferase reporter, and the HKU1 spike plasmid. Pseudotyped virions were harvested 2 and 3 days after transfection. Production efficacy was assessed by measuring infectivity or HIV Gag p24 concentration. For infection assays, 293T cells (6 × 10^4^) were transfected in suspension with Lipofectamine 2000 (Thermo) in a 96-well white plate (20ng of TMPRSS2 plasmids adjusted to 100 ng DNA with pQCXIP-Empty). 24 h post-transfection, cells were passed in 2 wells, and infected with indicated amount of virus (5-10 ng of p24) in 100 µL. The next day, 100 µL of media was added. 48 h post-infection, 125 µL of media was carefully removed, and 75 µL of Bright-Glo™ lysis buffer (ProMega) was added. After 10 min, luminescence was acquired using the EnSpire in a white plate (PerkinElmer).

### Flow cytometry

For spike binding, 293T cells were transiently transfected with TMPRSS2 and incubated with Camostat (10 μM) for 2 h. The cells were incubated with soluble biotinylated spike diluted in MACS buffer (PBS, 5 g/L BSA, 2 mM EDTA) at 2 µg/mL for 30 min at 4°C. The cells were then washed twice with PBS and then incubated with Alexa Fluor 647-conjugated streptavidin (Thermo Fisher Scientific, S21374, 1:400) for 30 min at 4°C.

For the spike, transfection efficiency was measured at the surface of live cells using mAb10 diluted in MACS buffer for 30 min at 4°C, and a human secondary IgG. mAb10 is an antibody generated from a SARS-CoV-2 infected patient which cross-reacts with HKU1 ^53^.

Surface expression of TMPRSS2 was assessed on live cells by staining with anti-TMPRSS2 A01-Fc ^1^ at 1 µg/mL, for 30 min at 4°C in MACS buffer, followed by staining with Alexa Fluor 647-conjugated Goat anti-Human antibody (Thermo Fisher Scientific, A-21445, 1:500).

TMPRSS2-myc expression was assessed on fixed cells by staining intracellularly with anti-cMyc 9E10 (Thermo -M4439, 1:400), for 30 min at RT in PBS/BSA/Azide/0.05% Saponin followed by Alexa Fluor 647 Goat anti-mouse antibody (Thermo Fisher Scientific, A-21242, 1:500).

All cells were washed twice with PBS and fixed with 4% paraformaldehyde. The results were acquired using an Attune Nxt Flow Cytometer (Life Technologies, software v3.2.1). Gating strategies are described in Figure SI 7.

## DATA AVAILABILITY

The data generated in this study and related to the X-ray structures determined have been deposited to the RCSB protein databank under PDB accession codes 8S0M (ternary complex RBD/TMPRSS^S441A^/A01), 8S0L (complex TMPRSS2^S441A^/A07, crystal form 1, higher resolution dataset), 8S0N (complex TMPRSS2^S441A^/A07, crystal form 2).

## ACKNOWLEDGEMENT

We thank the staff from the Crystallography platform at Institut Pasteur and the synchrotron source SOLEIL (Saint-Aubin, France) for granting access to the facility. We thank the staff of the beamlines Proxima 1, especially Pierre Legrand, and Proxima 2A for their advice and assistance during X-ray data collections. We thank Pablo Guardado-Calvo for granting access to the Octet-R8 instrument and Patrick England from the Molecular Biophysics platform at Institut Pasteur for useful discussions on biolayer interferometry data analysis.

This work was supported by the ‘URGENCE COVID-19’ fundraising campaign of Institut Pasteur. F.A.R. is funded by ANR grant nos. ANR-13-ISV8-0002-01 and ANR-10-LABX-62-10 IBEID, Wellcome Trust collaborative grant no. UNS22082 and the Institut Pasteur and the Centre national de la recherche scientifique. O.S. is funded by Institut Pasteur, Fondation pour la Recherche Médicale, the National Agency for AIDS Research–Emerging Infectious Diseases, the Vaccine Research Institute (ANR-10-LABX-77), Humanities in the European Research Area programme (DURABLE consortium), Agence Nationale de la Recherche (ANR)/Fondation pour la Recherche Médicale Flash Covid PROTEO-SARS-CoV-2 and IDISCOVR, and IBEID Labex. Work with UtechS Photonic BioImaging is funded by grant no. ANR-10-INSB-04-01 and Région Ile-de-France programme DIM1-Health

## Contributions

The experimental strategy was designed by I.F, N.S., S.D., J.B., O.S. and F.A.R. Construct design, protein expression and protein purification were performed by I.F., A.A and E.B.S. Crystallization and X-ray diffraction data collection were carried out by I.F. and A.H., while data processing was performed by I.F. and S.D. Cell-cell fusion, spike binding and pseudovirus entry experiments, as well as the mutagenesis of the plasmids used for these assays were performed by N.S., W.H.B. and J.B. The biolayer interferometry experiments were performed by I.F. The original manuscript draft was written by I.F. and F.A.R., with input from N.S., S.D., J.B. and O.S. The funding was acquired by O.S. and F.A.R.

## Declaration of interests

I.F., N.S., E.B.S., P.L. J.B., O.S. and F.A.R. have a provisional patent on anti-TMPRSS2 nanobodies.

F.A.R. is a founder of *Meletios Therapeutics* and a member of its scientific advisory board. The other authors declare no competing interests.

## SUPPLEMENTARY INFORMATION

**Figure SI-1.**
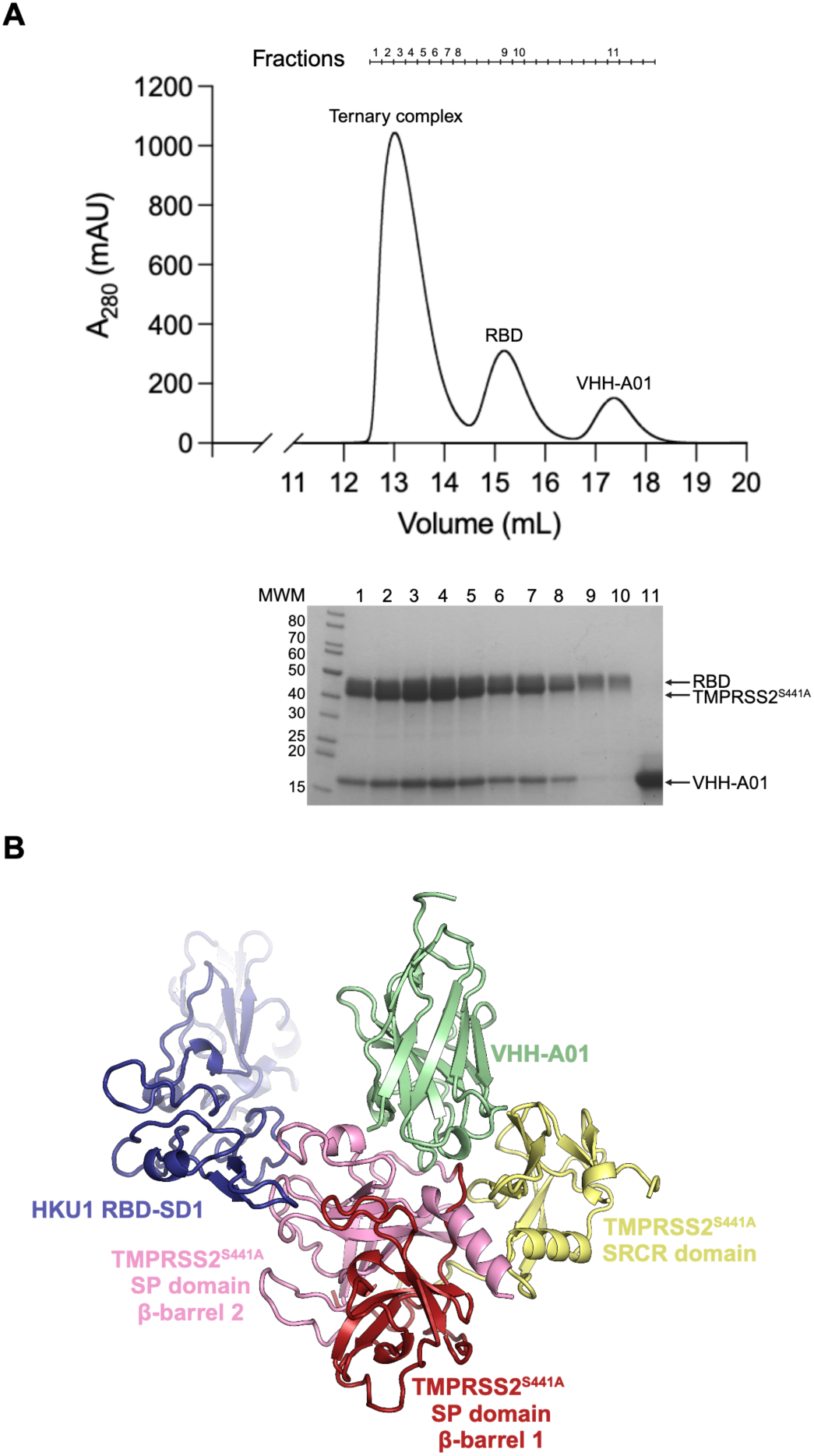
(Related to Figure 1) Purification of the ternary complex TMPRSS2^S441A^/RBD-SD1/VHH-A01. (A) Size-exclusion chromatography of the binding reaction prepared with HKU1B RBD, TMPRSS2^S441A^ and VHH-A01. The eluate was collected in different fractions (numbers indicated above the chromatogram) and 10 μl aliquots of some of them were analyzed under reducing SDS-PAGE (bottom panel). MWM: molecular weight marker. (B) Crystal structure of the ternary complex.

**Figure SI-2.**
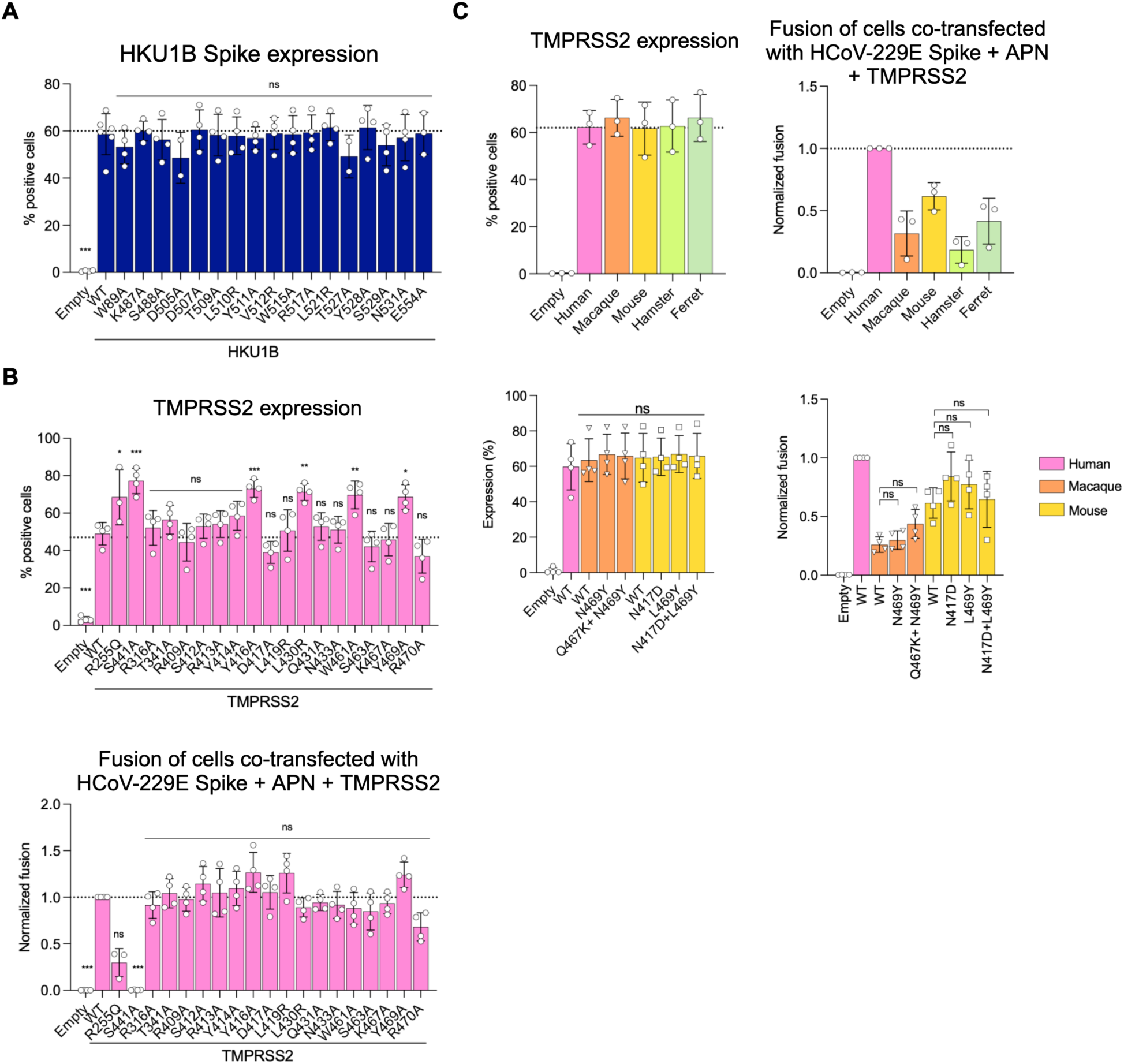
(Related to Figure 2). Expression and activity of Spike mutants, TMPRSS2 mutants and TMPRSS2 orthologs. (A) Expression of HKU1B Spike constructs determined by FACS using an anti-S2 antibody. (B) Surface expression of TMPRSS2 mutants assessed on live cells by FACS with anti-TMPRSS2 VHH-A01-Fc (top). Cell-cell fusion assay performed by co-transfecting cells with HCoV-229E Spike, APN (aminopeptidase N, the receptor) and a TMPRSS2 variant from the panel (bottom panel). (C) Expression of TMPRSS2 orthologs and their point mutants evaluated by intracellular staining of their c-myc epitope (left panel). Cell-cell fusion assay performed by co-transfecting cells with HCoV-229E Spike, APN and one TMPRSS2 ortholog or point mutant (right panel) Statistical analysis: (A, B, C) One-way ANOVA with Dunnett’s multiple comparison test compared to the WT. For fusion assay One-Way ANOVA was performed on the non-normalized log transformed data. * p<0.05, ** p<0.01, *** p<0.001.

**Figure SI-3.**
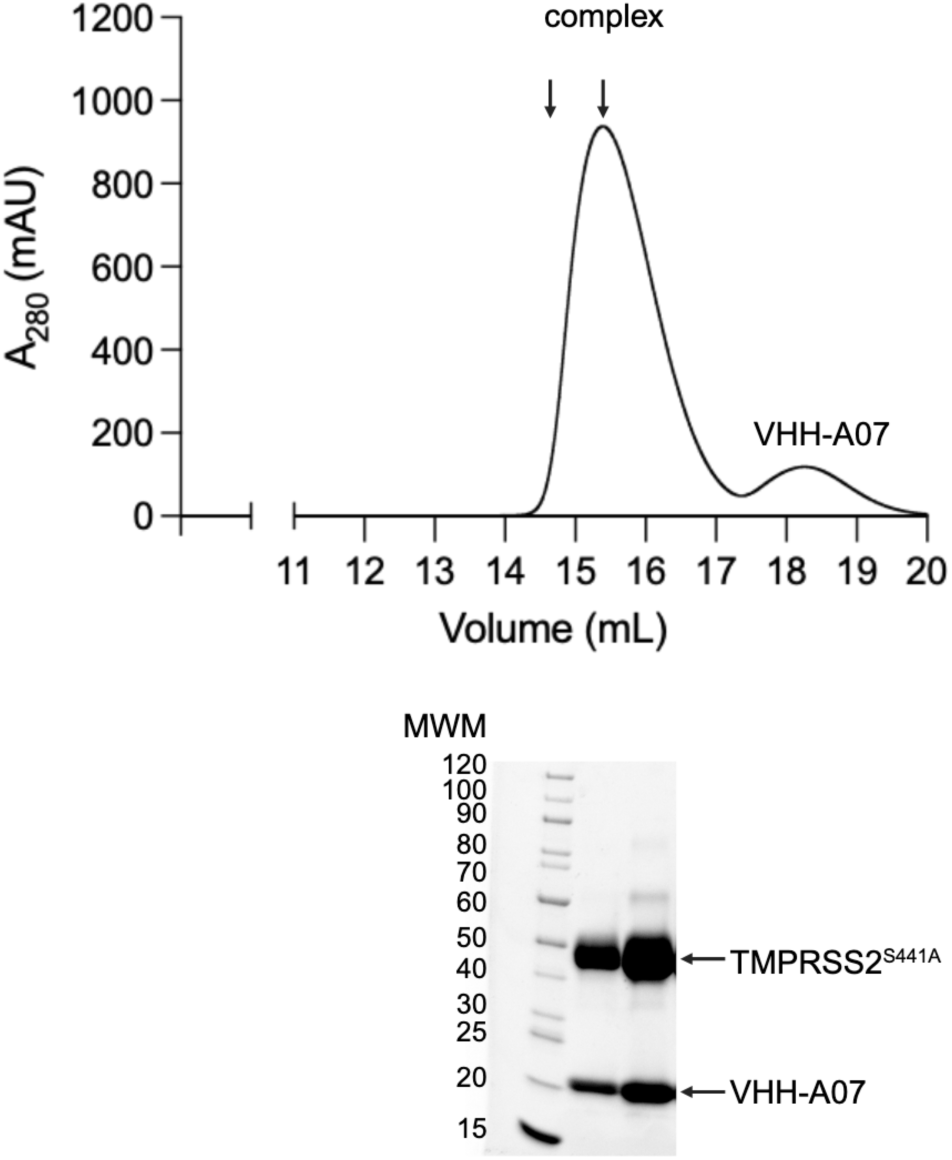
(Related to Figure 4). Purification of the TMPRSS2^S441A^+VHH-A07 complex. TMPRSS2^S441A^ was incubated with an excess of VHH-A07 and the binding reaction was injected onto a size-exclusion chromatography column. The elution profile is shown, with two arrows indicating fractions that were analyzed by reducing SDS-PAGE (bottom). MWM: molecular weight marker.

**Figure SI-4.**
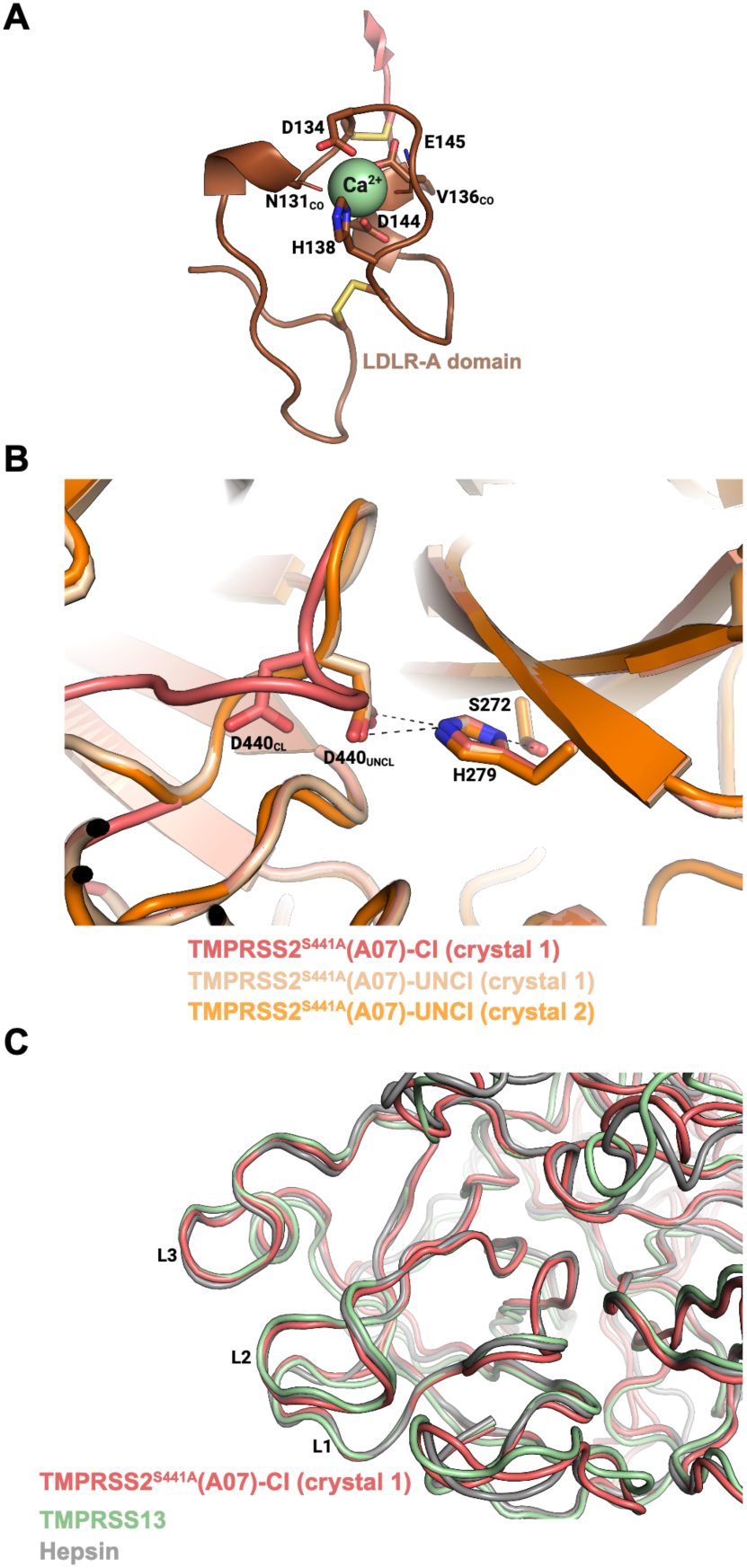
(Related to Figure 4). TMPRSS2^S441A^ structure. (A) Structure of the LDLR-A domain obtained from crystal form 1 of the TMPRSS2^S441A^+VHH-A07 complex. A central calcium ion is shown, as well as the two backbone carbonyls (subscript CO) and side chains that coordinate it. Disulfides C133-C148 and C120-C139 are shown with yellow sticks. (B) Superposition of the structures obtained for cleaved TMPRSS2^S441A^ in the crystal 1 (TMPRSS2^S441A^-Cl, red), TMPRSS2^S441A^ zymogen in crystal 1 (TMPRSS2^S441A^-UNCl, wheat) and TMPRSS2^S441A^ zymogen from crystal 2 (TMPRSS2^S441A^-UNCl, orange) showing the zymogen triad (S272, H279, D440). Upon autocleavage, the conformational change in loop 1 shifts the position of D440 (indicated by the subscript ‘Cl’). (C) Superposition of cleaved TMPRSS2^S441A^ (from crystal 1) with active proteases from its subfamily (TMPRSS13, green; hepsin, gray). The loops 1, 2 and 3 (L1, L2 and L3, respectively) are labeled.

**Figure SI-5.**
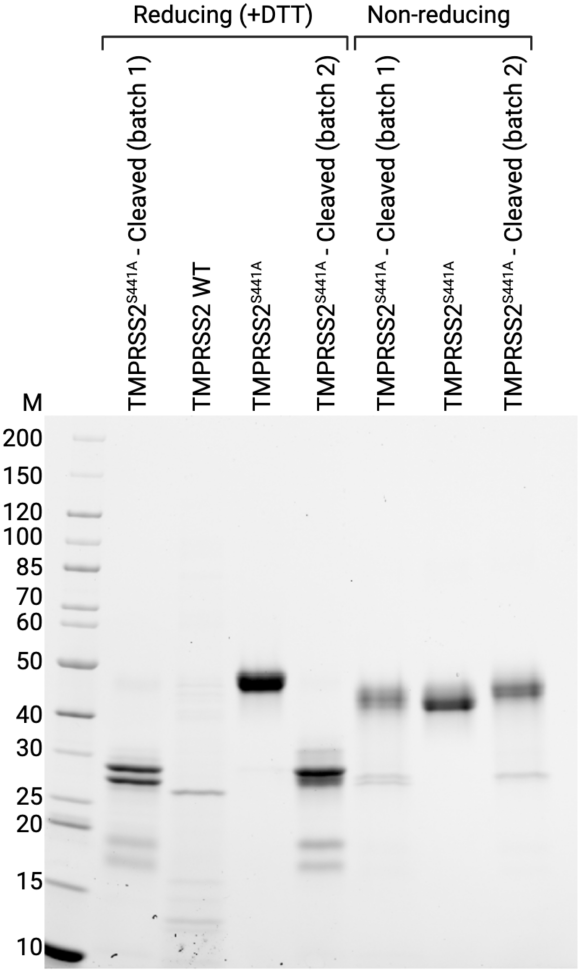
(Related to FIgure 4). TMPRSS2^S441A^ proteolysis by TMPRSS2 wild-type. SDS-PAGE under reducing and non-reducing conditions to show the cleavage of TMPRSS2^S441A^ upon incubation with the wild-type protease. Two reactions were performed under different conditions (producing ‘batch 1’and ‘batch 2’) and 2 μl of each reaction (corresponding to ∼2.5 μg of substrate) were analyzed by SDS-PAGE. For comparison, wild-type (WT) TMPRSS2 and untreated TMPRSS2^S441A^ were also loaded in the gel. M: molecular weight marker.

**Figure SI-6.**
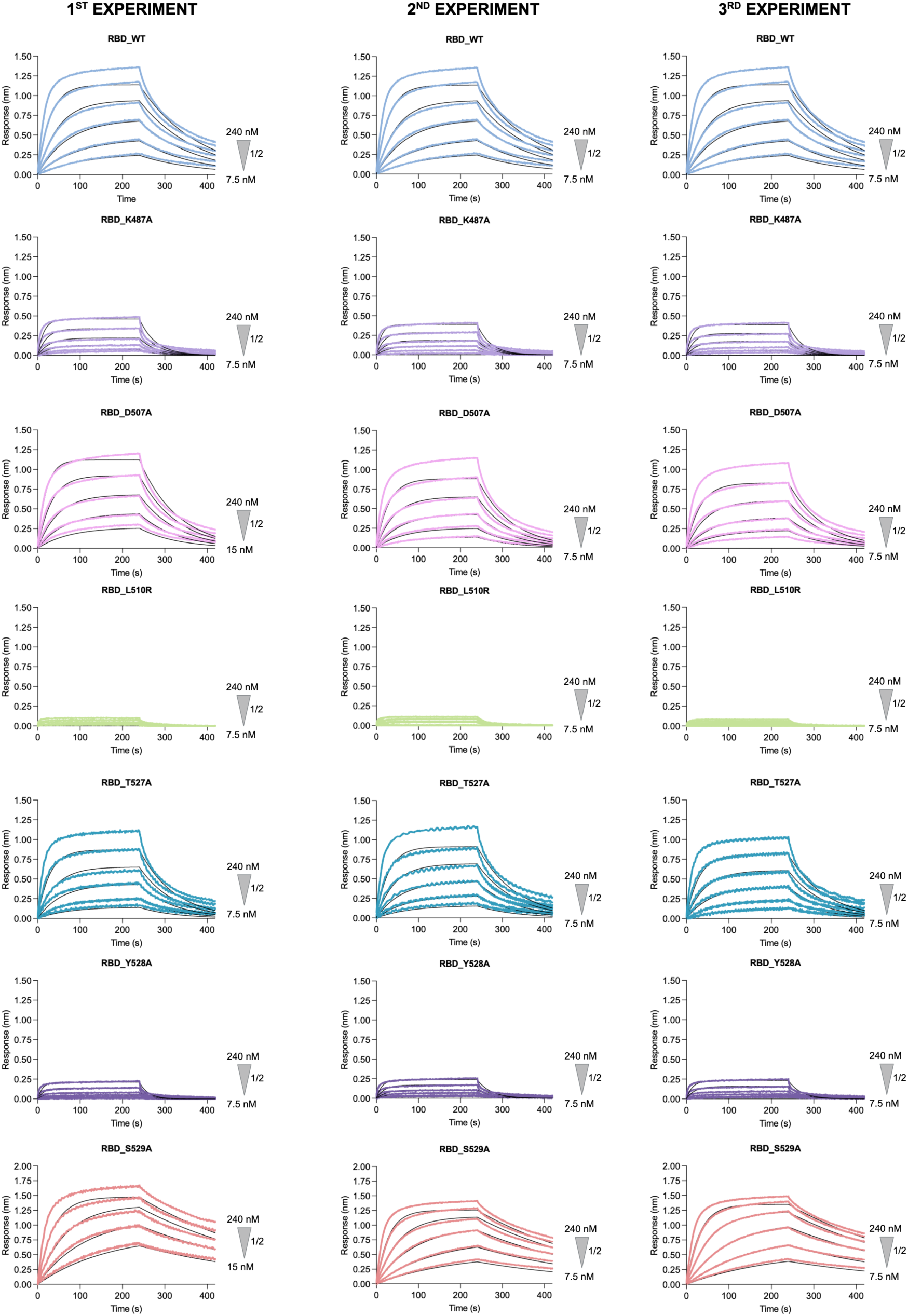
(Related to Figure 5). Biolayer interferometry (BLI) experiments to determine the Kd of RBD mutants against cleaved TMPRSS2^S441A^.

**Figure SI-7.**
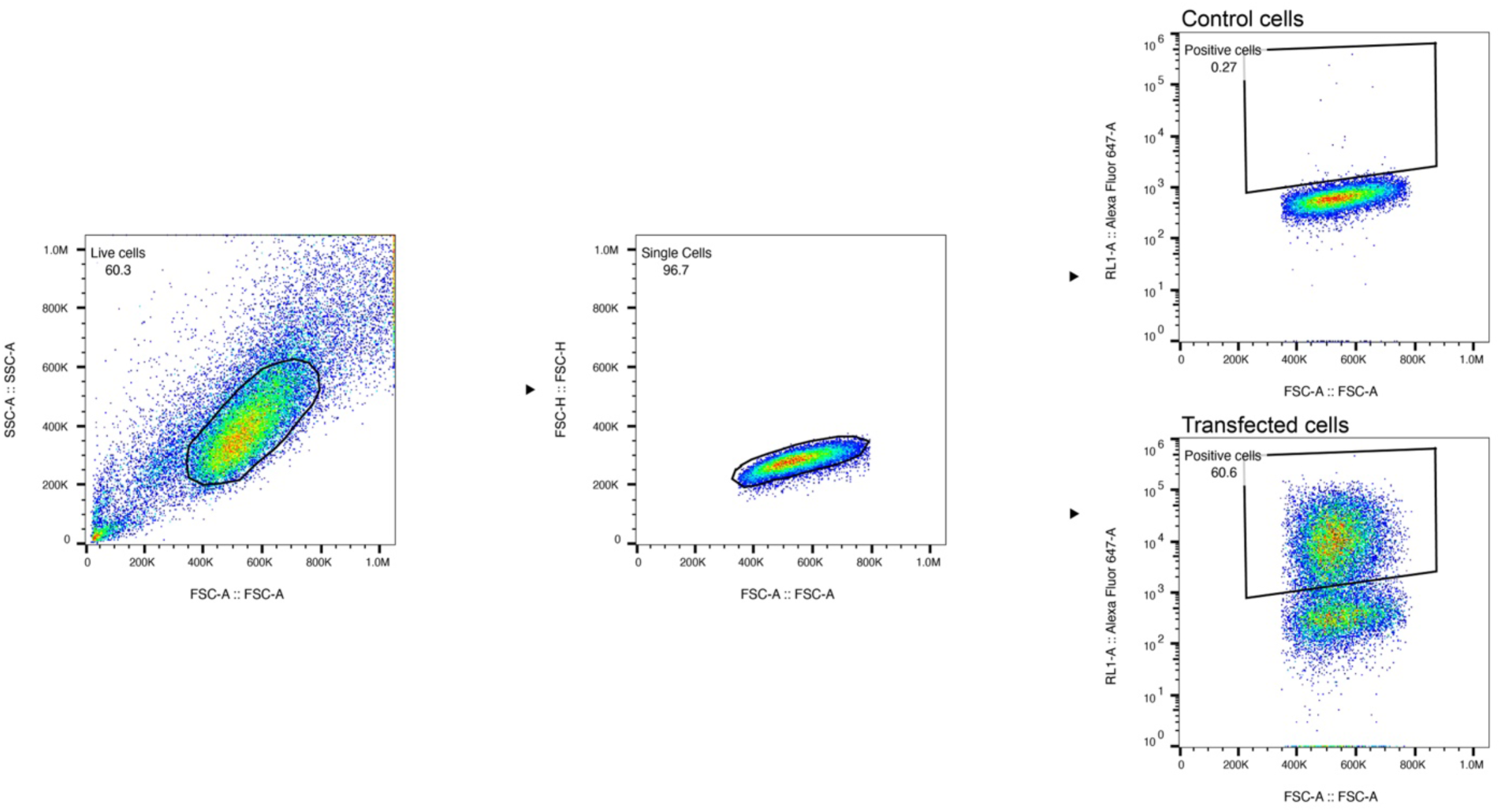
(Related to Methods and to FIgure 2). Gating strategy for the detection of surface expression of the transfected proteins by flow cytometry.

**Table SI1.**
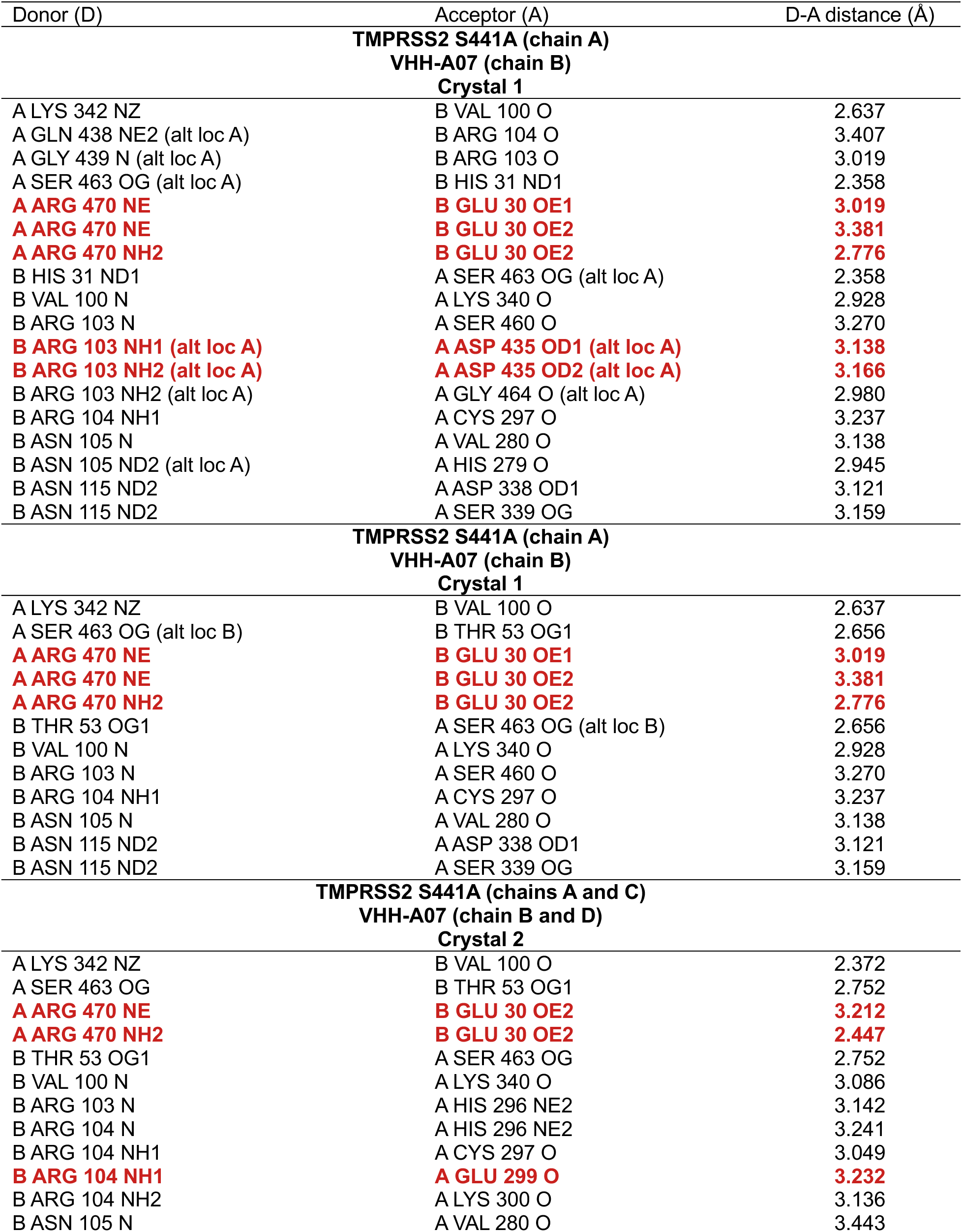
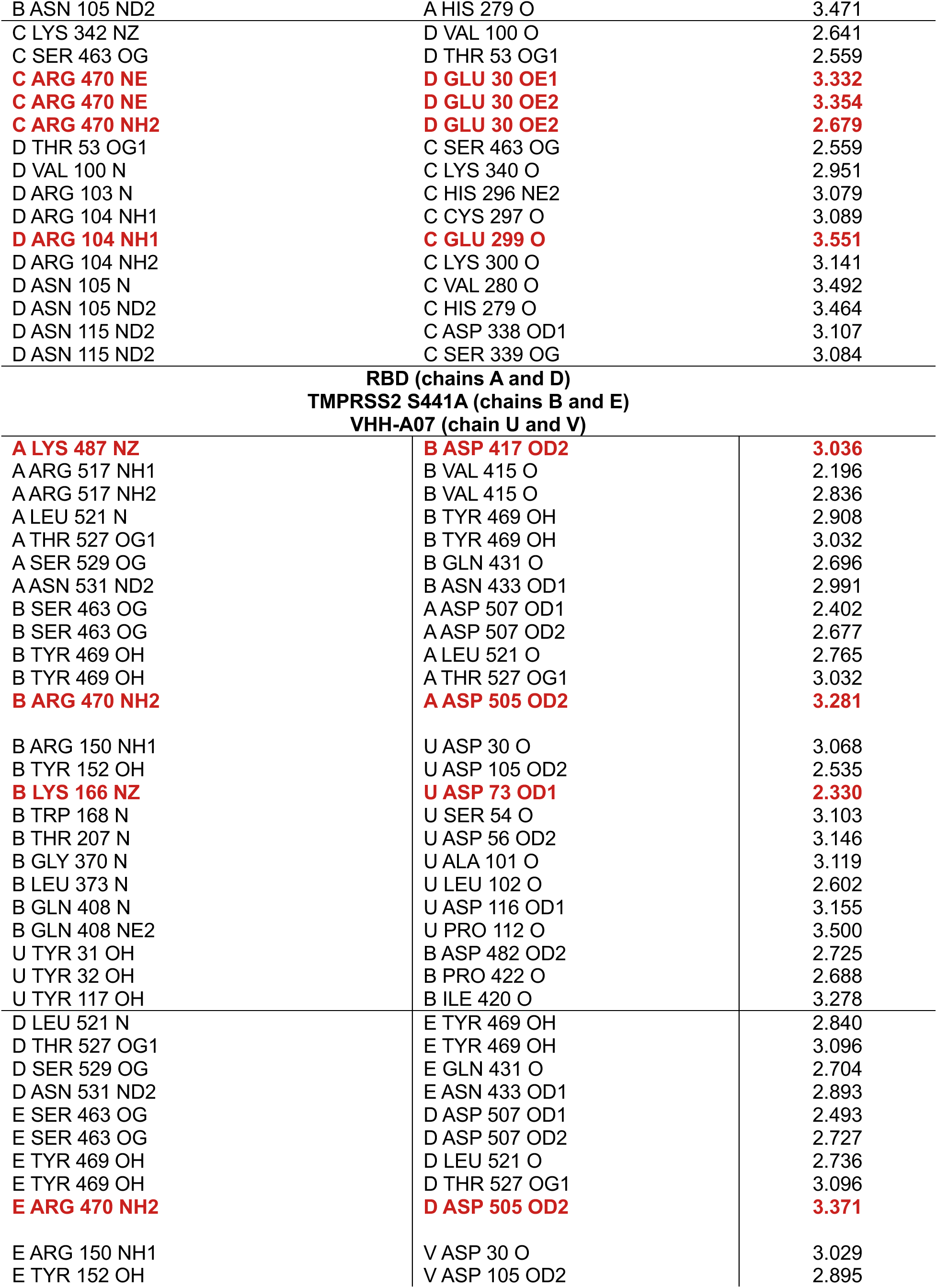
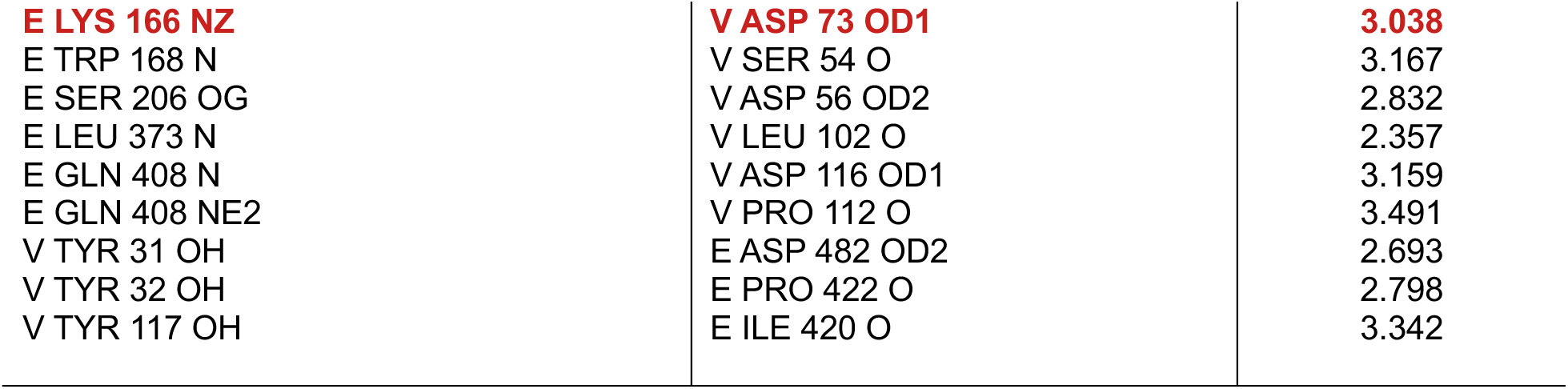
(Related to FIgure 1). List of residues involved in hydrogen bonds and salt bridges.

## REFERENCES

1. Saunders, N., Fernandez, I., Planchais, C., Michel, V., Rajah, M.M., Baquero Salazar, E., Postal, J., Porrot, F., Guivel-Benhassine, F., Blanc, C., et al. (2023). TMPRSS2 is a functional receptor for human coronavirus HKU1. Nature 624, 207–214. 10.1038/s41586-023-06761-7.

2. Ko, C.J., Huang, C.C., Lin, H.Y., Juan, C.P., Lan, S.W., Shyu, H.Y., Wu, S.R., Hsiao, P.W., Huang, H.P., Shun, C.T., and Lee, M.S. (2015). Androgen-Induced TMPRSS2 Activates Matriptase and Promotes Extracellular Matrix Degradation, Prostate Cancer Cell Invasion, Tumor Growth, and Metastasis. Cancer Res 75, 2949–2960. 10.1158/0008-5472.CAN-14-3297.

3. Mukai, S., Yorita, K., Kawagoe, Y., Katayama, Y., Nakahara, K., Kamibeppu, T., Sugie, S., Tukino, H., Kamoto, T., and Kataoka, H. (2015). Matriptase and MET are prominently expressed at the site of bone metastasis in renal cell carcinoma: immunohistochemical analysis. Hum Cell 28, 44–50. 10.1007/s13577-014-0101-3.

4. Tomlins, S.A., Rhodes, D.R., Perner, S., Dhanasekaran, S.M., Mehra, R., Sun, X.W., Varambally, S., Cao, X., Tchinda, J., Kuefer, R., et al. (2005). Recurrent fusion of TMPRSS2 and ETS transcription factor genes in prostate cancer. Science 310, 644–648. 10.1126/science.1117679.

5. Wang, Z., Wang, Y., Zhang, J., Hu, Q., Zhi, F., Zhang, S., Mao, D., Zhang, Y., and Liang, H. (2017). Significance of the TMPRSS2:ERG gene fusion in prostate cancer. Mol Med Rep 16, 5450–5458. 10.3892/mmr.2017.7281.

6. Lucas, J.M., Heinlein, C., Kim, T., Hernandez, S.A., Malik, M.S., True, L.D., Morrissey, C., Corey, E., Montgomery, B., Mostaghel, E., et al. (2014). The androgen-regulated protease TMPRSS2 activates a proteolytic cascade involving components of the tumor microenvironment and promotes prostate cancer metastasis. Cancer Discov 4, 1310–1325. 10.1158/2159-8290.CD-13-1010.

7. Kahn, J.S., and McIntosh, K. (2005). History and recent advances in coronavirus discovery. Pediatr Infect Dis J 24, S223–227, discussion S226. 10.1097/01.inf.0000188166.17324.60.

8. Tortorici, M.A., and Veesler, D. (2019). Structural insights into coronavirus entry. Adv Virus Res 105, 93–116. 10.1016/bs.aivir.2019.08.002.

9. Huang, X., Dong, W., Milewska, A., Golda, A., Qi, Y., Zhu, Q.K., Marasco, W.A., Baric, R.S., Sims, A.C., Pyrc, K., et al. (2015). Human Coronavirus HKU1 Spike Protein Uses O-Acetylated Sialic Acid as an Attachment Receptor Determinant and Employs Hemagglutinin-Esterase Protein as a Receptor-Destroying Enzyme. J Virol 89, 7202–7213. 10.1128/JVI.00854-15.

10. Hulswit, R.J.G., Lang, Y., Bakkers, M.J.G., Li, W., Li, Z., Schouten, A., Ophorst, B., van Kuppeveld, F.J.M., Boons, G.J., Bosch, B.J., et al. (2019). Human coronaviruses OC43 and HKU1 bind to 9-O-acetylated sialic acids via a conserved receptor-binding site in spike protein domain A. Proc Natl Acad Sci U S A 116, 2681–2690. 10.1073/pnas.1809667116.

11. Li, Z., Lang, Y., Liu, L., Bunyatov, M.I., Sarmiento, A.I., de Groot, R.J., and Boons, G.J. (2021). Synthetic O-acetylated sialosides facilitate functional receptor identification for human respiratory viruses. Nat Chem 13, 496–503. 10.1038/s41557-021-00655-9.

12. Pronker, M.F., Creutznacher, R., Drulyte, I., Hulswit, R.J.G., Li, Z., van Kuppeveld, F.J.M., Snijder, J., Lang, Y., Bosch, B.J., Boons, G.J., et al. (2023). Sialoglycan binding triggers spike opening in a human coronavirus. Nature 624, 201–206. 10.1038/s41586-023-06599-z.

13. Benton, D.J., Wrobel, A.G., Xu, P., Roustan, C., Martin, S.R., Rosenthal, P.B., Skehel, J.J., and Gamblin, S.J. (2020). Receptor binding and priming of the spike protein of SARS-CoV-2 for membrane fusion. Nature 588, 327–330. 10.1038/s41586-020-2772-0.

14. Walls, A.C., Xiong, X., Park, Y.J., Tortorici, M.A., Snijder, J., Quispe, J., Cameroni, E., Gopal, R., Dai, M., Lanzavecchia, A., et al. (2020). Unexpected Receptor Functional Mimicry Elucidates Activation of Coronavirus Fusion. Cell 183, 1732. 10.1016/j.cell.2020.11.031.

15. Chen, Y.W., Lee, M.S., Lucht, A., Chou, F.P., Huang, W., Havighurst, T.C., Kim, K., Wang, J.K., Antalis, T.M., Johnson, M.D., and Lin, C.Y. (2010). TMPRSS2, a serine protease expressed in the prostate on the apical surface of luminal epithelial cells and released into semen in prostasomes, is misregulated in prostate cancer cells. Am J Pathol 176, 2986–2996. 10.2353/ajpath.2010.090665.

16. Afar, D.E., Vivanco, I., Hubert, R.S., Kuo, J., Chen, E., Saffran, D.C., Raitano, A.B., and Jakobovits, A. (2001). Catalytic cleavage of the androgen-regulated TMPRSS2 protease results in its secretion by prostate and prostate cancer epithelia. Cancer Res 61, 1686–1692.

17. Tseng, C.C., Jia, B., Barndt, R., Gu, Y., Chen, C.Y., Tseng, I.C., Su, S.F., Wang, J.K., Johnson, M.D., and Lin, C.Y. (2017). Matriptase shedding is closely coupled with matriptase zymogen activation and requires de novo proteolytic cleavage likely involving its own activity. PLoS One 12, e0183507. 10.1371/journal.pone.0183507.

18. Fraser, B.J., Beldar, S., Seitova, A., Hutchinson, A., Mannar, D., Li, Y., Kwon, D., Tan, R., Wilson, R.P., Leopold, K., et al. (2022). Structure and activity of human TMPRSS2 protease implicated in SARS-CoV-2 activation. Nat Chem Biol 18, 963–971. 10.1038/s41589-022-01059-7.

19. Bertram, S., Dijkman, R., Habjan, M., Heurich, A., Gierer, S., Glowacka, I., Welsch, K., Winkler, M., Schneider, H., Hofmann-Winkler, H., et al. (2013). TMPRSS2 activates the human coronavirus 229E for cathepsin-independent host cell entry and is expressed in viral target cells in the respiratory epithelium. J Virol 87, 6150–6160. 10.1128/JVI.03372-12.

20. Glowacka, I., Bertram, S., Muller, M.A., Allen, P., Soilleux, E., Pfefferle, S., Steffen, I., Tsegaye, T.S., He, Y., Gnirss, K., et al. (2011). Evidence that TMPRSS2 activates the severe acute respiratory syndrome coronavirus spike protein for membrane fusion and reduces viral control by the humoral immune response. J Virol 85, 4122–4134. 10.1128/JVI.02232-10.

21. Hoffmann, M., Kleine-Weber, H., Schroeder, S., Kruger, N., Herrler, T., Erichsen, S., Schiergens, T.S., Herrler, G., Wu, N.H., Nitsche, A., et al. (2020). SARS-CoV-2 Cell Entry Depends on ACE2 and TMPRSS2 and Is Blocked by a Clinically Proven Protease Inhibitor. Cell 181, 271–280 e278. 10.1016/j.cell.2020.02.052.

22. Limburg, H., Harbig, A., Bestle, D., Stein, D.A., Moulton, H.M., Jaeger, J., Janga, H., Hardes, K., Koepke, J., Schulte, L., et al. (2019). TMPRSS2 Is the Major Activating Protease of Influenza A Virus in Primary Human Airway Cells and Influenza B Virus in Human Type II Pneumocytes. J Virol 93. 10.1128/JVI.00649-19.

23. Shirato, K., Kawase, M., and Matsuyama, S. (2013). Middle East respiratory syndrome coronavirus infection mediated by the transmembrane serine protease TMPRSS2. J Virol 87, 12552–12561. 10.1128/JVI.01890-13.

24. Perona, J.J., and Craik, C.S. (1995). Structural basis of substrate specificity in the serine proteases. Protein Sci 4, 337–360. 10.1002/pro.5560040301.

25. Goettig, P., Brandstetter, H., and Magdolen, V. (2019). Surface loops of trypsin-like serine proteases as determinants of function. Biochimie 166, 52–76. 10.1016/j.biochi.2019.09.004.

26. Perona, J.J., and Craik, C.S. (1997). Evolutionary divergence of substrate specificity within the chymotrypsin-like serine protease fold. J Biol Chem 272, 29987–29990. 10.1074/jbc.272.48.29987.

27. Lawrence, M.C., and Colman, P.M. (1993). Shape complementarity at protein/protein interfaces. J Mol Biol 234, 946–950. 10.1006/jmbi.1993.1648.

28. Kirchdoerfer, R.N., Cottrell, C.A., Wang, N., Pallesen, J., Yassine, H.M., Turner, H.L., Corbett, K.S., Graham, B.S., McLellan, J.S., and Ward, A.B. (2016). Pre-fusion structure of a human coronavirus spike protein. Nature 531, 118–121. 10.1038/nature17200.

29. Yeager, C.L., Ashmun, R.A., Williams, R.K., Cardellichio, C.B., Shapiro, L.H., Look, A.T., and Holmes, K.V. (1992). Human aminopeptidase N is a receptor for human coronavirus 229E. Nature 357, 420–422. 10.1038/357420a0.

30. Pasternak, A., Ringe, D., and Hedstrom, L. (1999). Comparison of anionic and cationic trypsinogens: the anionic activation domain is more flexible in solution and differs in its mode of BPTI binding in the crystal structure. Protein Sci 8, 253–258. 10.1110/ps.8.1.253.

31. Freer, S.T., Kraut, J., Robertus, J.D., Wright, H.T., and Xuong, N.H. (1970). Chymotrypsinogen: 2.5-angstrom crystal structure, comparison with alpha-chymotrypsin, and implications for zymogen activation. Biochemistry 9, 1997–2009. 10.1021/bi00811a022.

32. Madison, E.L., Kobe, A., Gething, M.J., Sambrook, J.F., and Goldsmith, E.J. (1993). Converting tissue plasminogen activator to a zymogen: a regulatory triad of Asp-His-Ser. Science 262, 419–421. 10.1126/science.8211162.

33. Herter, S., Piper, D.E., Aaron, W., Gabriele, T., Cutler, G., Cao, P., Bhatt, A.S., Choe, Y., Craik, C.S., Walker, N., et al. (2005). Hepatocyte growth factor is a preferred in vitro substrate for human hepsin, a membrane-anchored serine protease implicated in prostate and ovarian cancers. Biochem J 390, 125–136. 10.1042/BJ20041955.

34. Ohno, A., Maita, N., Tabata, T., Nagano, H., Arita, K., Ariyoshi, M., Uchida, T., Nakao, R., Ulla, A., Sugiura, K., et al. (2021). Crystal structure of inhibitor-bound human MSPL that can activate high pathogenic avian influenza. Life Sci Alliance 4. 10.26508/lsa.202000849.

35. McCallum, M., Park, Y.J., Stewart, C., Sprouse, K.R., Brown, J., Tortorici, M.A., Gibson, C., Wong, E., Ieven, M., Telenti, A., and Veesler, D. (2024). Human coronavirus HKU1 recognition of the TMPRSS2 host receptor. bioRxiv. 10.1101/2024.01.09.574565.

36. Ke, Z., Oton, J., Qu, K., Cortese, M., Zila, V., McKeane, L., Nakane, T., Zivanov, J., Neufeldt, C.J., Cerikan, B., et al. (2020). Structures and distributions of SARS-CoV-2 spike proteins on intact virions. Nature 588, 498–502. 10.1038/s41586-020-2665-2.

37. Kerr, M.A., Walsh, K.A., and Neurath, H. (1976). A proposal for the mechanism of chymotrypsinogen activation. Biochemistry 15, 5566–5570. 10.1021/bi00670a022.

38. Buchrieser, J., Dufloo, J., Hubert, M., Monel, B., Planas, D., Rajah, M.M., Planchais, C., Porrot, F., Guivel-Benhassine, F., Van der Werf, S., et al. (2020). Syncytia formation by SARS-CoV-2-infected cells. EMBO J 39, e106267. 10.15252/embj.2020106267.

39. Kodaka, M., Yang, Z., Nakagawa, K., Maruyama, J., Xu, X., Sarkar, A., Ichimura, A., Nasu, Y., Ozawa, T., Iwasa, H., et al. (2015). A new cell-based assay to evaluate myogenesis in mouse myoblast C2C12 cells. Exp Cell Res 336, 171–181. 10.1016/j.yexcr.2015.06.015.

40. Edie, S., Zaghloul, N.A., Leitch, C.C., Klinedinst, D.K., Lebron, J., Thole, J.F., McCallion, A.S., Katsanis, N., and Reeves, R.H. (2018). Survey of Human Chromosome 21 Gene Expression Effects on Early Development in Danio rerio. G3 (Bethesda) 8, 2215–2223. 10.1534/g3.118.200144.

41. Hoffmann, M., Hofmann-Winkler, H., Smith, J.C., Kruger, N., Arora, P., Sorensen, L.K., Sogaard, O.S., Hasselstrom, J.B., Winkler, M., Hempel, T., et al. (2021). Camostat mesylate inhibits SARS-CoV-2 activation by TMPRSS2-related proteases and its metabolite GBPA exerts antiviral activity. EBioMedicine 65, 103255. 10.1016/j.ebiom.2021.103255.

42. Chavas, L.M.G., Gourhant, P., Guimaraes, B.G., Isabet, T., Legrand, P., Lener, R., Montaville, P., Sirigu, S., and Thompson, A. (2021). PROXIMA-1 beamline for macromolecular crystallography measurements at Synchrotron SOLEIL. J Synchrotron Radiat 28, 970–976. 10.1107/S1600577521002605.

43. Kabsch, W. (2010). Integration, scaling, space-group assignment and post-refinement. Acta Crystallogr D Biol Crystallogr 66, 133–144. 10.1107/S0907444909047374.

44. Kabsch, W. (2010). Xds. Acta Crystallogr D Biol Crystallogr 66, 125–132. 10.1107/S0907444909047337.

45. Evans, P.R., and Murshudov, G.N. (2013). How good are my data and what is the resolution? Acta Crystallogr D Biol Crystallogr 69, 1204–1214. 10.1107/S0907444913000061.

46. Liebschner, D., Afonine, P.V., Baker, M.L., Bunkoczi, G., Chen, V.B., Croll, T.I., Hintze, B., Hung, L.W., Jain, S., McCoy, A.J., et al. (2019). Macromolecular structure determination using X-rays, neutrons and electrons: recent developments in Phenix. Acta Crystallogr D Struct Biol 75, 861–877. 10.1107/S2059798319011471.

47. Emsley, P., Lohkamp, B., Scott, W.G., and Cowtan, K. (2010). Features and development of Coot. Acta Crystallogr D Biol Crystallogr 66, 486–501. 10.1107/S0907444910007493.

48. Williams, C.J., Headd, J.J., Moriarty, N.W., Prisant, M.G., Videau, L.L., Deis, L.N., Verma, V., Keedy, D.A., Hintze, B.J., Chen, V.B., et al. (2018). MolProbity: More and better reference data for improved all-atom structure validation. Protein Sci 27, 293–315. 10.1002/pro.3330.

49. Krissinel, E., and Henrick, K. (2007). Inference of macromolecular assemblies from crystalline state. J Mol Biol 372, 774–797. 10.1016/j.jmb.2007.05.022.

50. DeLano, W.L. (2002). The PyMOL Molecular Graphics System.. DeLano Scientific, San Carlos, CA, USA, 2002.

51. Madeira, F., Pearce, M., Tivey, A.R.N., Basutkar, P., Lee, J., Edbali, O., Madhusoodanan, N., Kolesnikov, A., and Lopez, R. (2022). Search and sequence analysis tools services from EMBL-EBI in 2022. Nucleic Acids Res 50, W276–W279. 10.1093/nar/gkac240.

52. Crooks, G.E., Hon, G., Chandonia, J.M., and Brenner, S.E. (2004). WebLogo: a sequence logo generator. Genome Res 14, 1188–1190. 10.1101/gr.849004.

53. Planchais, C., Fernandez, I., Bruel, T., de Melo, G.D., Prot, M., Beretta, M., Guardado-Calvo, P., Dufloo, J., Molinos-Albert, L.M., Backovic, M., et al. (2022). Potent human broadly SARS-CoV-2-neutralizing IgA and IgG antibodies effective against Omicron BA.1 and BA.2. J Exp Med 219. 10.1084/jem.20220638.

